# MND1 and PSMC3IP control PARP inhibitor sensitivity in mitotic cells

**DOI:** 10.1101/2022.08.24.505108

**Authors:** Anabel Zelceski, Paola Francica, Lea Lingg, Merve Mutlu, Colin Stok, Martin Liptay, John Alexander, Joseph S. Baxter, Rachel Brough, Aditi Gulati, Syed Haider, Maya Raghunandan, Feifei Song, Sandhya Sridhar, Josep V. Forment, Mark J. O’Connor, Barry R. Davies, Marcel A.T.M. van Vugt, Dragomir B. Krastev, Stephen J. Pettitt, Andrew N. J. Tutt, Sven Rottenberg, Christopher J. Lord

**Affiliations:** The CRUK Gene Function Laboratory and The Institute of Cancer Research, London, SW3 6JB, UK; Breast Cancer Now Toby Robins Breast Cancer Research Centre, The Institute of Cancer Research, London, SW3 6JB, UK; Institute of Animal Pathology, Vetsuisse Faculty, University of Bern, 3012 Bern, Switzerland; Departement of Biomedical Research (DBMR), Cancer Therapy Resistance Cluster, University of Bern, 3012 Bern, Switzerland; Department of Medical Oncology, University Medical Center Groningen, University of Groningen, Hanzeplein 1, 9713GZ, Groningen, The Netherlands; Oncology R&D, AstraZeneca, Cambridge, United Kingdom; Division of Molecular Pathology, The Netherlands Cancer Institute, 1066CX Amsterdam, The Netherlands; Bern Center for Precision Medicine, University of Bern, 3012 Bern, Switzerland

**Keywords:** MND1, PSMC3IP, PARP inhibitor, DNA repair

## Abstract

The PSMC3IP-MND1 heterodimer promotes RAD51 and DMC1-dependent D-loop formation during meiosis in yeast and mammalian organisms. For this purpose, it catalyzes the DNA strand exchange activities of the recombinases. Interestingly, in a panel of genome-scale CRISPR-Cas9 mutagenesis and interference screens in mitotic cells, we found that depletion of either *PSMC3IP* or *MND1* caused sensitivity to clinical Poly (ADP-Ribose) Polymerase inhibitors (PARPi). A retroviral mutagenesis screen in mitotic cells also identified *PSMC3IP* and *MND1* as genetic determinants of ionizing radiation sensitivity. The role *PSMC3IP* and *MND1* play in preventing PARPi sensitivity in mitotic cells appears to be independent of a previously described role in alternative lengthening of telomeres (ALT). *PSMC3IP* or *MND1* depleted cells accumulate toxic RAD51 foci in response to DNA damage, show impaired homology-directed DNA repair, and become PARPi sensitive, even in cells lacking both *BRCA1* and *TP53BP1*. Although replication fork reversal is also affected, the epistatic relationship between *PSMC3IP-MND1* and *BRCA1/BRCA2* suggests that the abrogated D-loop formation is the major cause of PARPi sensitivity. This is corroborated by the fact that a *PSMC3IP* p.Glu201del D-loop formation mutant associated with ovarian dysgenesis fails to reverse PARPi sensitivity. These observations suggest that meiotic proteins such as MND1 and PSMC3IP could have a greater role in mitotic cells in determining the response to therapeutic DNA damage.

## Introduction

To date, five different Poly(ADP-Ribose) Polymerase inhibitors (PARPi) (Bryant et al., 2005; Farmer et al., 2005) have been approved for the treatment of homologous recombination (HR) defective cancers (Litton et al., 2018; Robson et al., 2017). Adapting the genetic concept of synthetic lethality to cancer therapy (Kaelin, 2005), PARPi are thought to work by generating a DNA lesion, most likely “trapping” of the DNA repair protein Poly(ADP-Ribose) Polymerase 1 (PARP1) (Murai et al., 2012) on chromatin at sites of damage. The nucleoprotein complex caused by PARP1 trapping provides a steric barrier to the normal function of DNA and impairs the normal progression of the replication fork (RF) (Krastev et al., 2021). The DNA damage caused by PARPi is normally repaired by BRCA1, BRCA2 and RAD51-mediated HR; in the absence of BRCA1 or BRCA2 function, PARPi-mediated cytotoxicity ensues (Farmer et al., 2005). Recent evidence also suggests that trapped PARP1 at replication gaps between Okazaki fragments is a major cause of PARPi sensitivity (Cong et al., 2021; Hanzlikova et al., 2018; Vaitsiankova et al., 2022). Targeting PARP1 in patients with HR-defective cancers is therefore a key example of the concept of precision medicine in cancer therapy (Drean et al., 2017).

To identify patients with HR-defective tumors, various approaches are used. This includes the detection of mutations in *BRCA1/2* or other HR-associated genes (Cancer Genome Atlas Research, 2011), prior platinum sensitivity (Ledermann et al., 2012), or a genomic mutational signature which reflects the lack of HR and predominance of other DNA repair pathways (Davies et al., 2017; Gulhan et al., 2019; Nik-Zainal et al., 2016; Polak et al., 2017). Experimentally, PARPi sensitivity can also be predicted by the inability to localize the DNA recombinase RAD51 to the site of DNA damage, an effect estimated by the absence of nuclear RAD51 foci (Cruz et al., 2018; Llop-Guevara et al., 2021; van Wijk et al., 2020).

In order to better understand what determines PARPi sensitivity, we carried out a series of parallel CRISPR mutagenesis and interference screens. Both mutagenesis and interference screens identified the meiotic recombination heterodimer MND1-PSMC3IP as controlling PARPi response in mitotic cells. Subsequent experiments showed that, in contrast to *BRCA1* or *BRCA2* mutant cells, *MND1-* or *PSMC3IP-* deficient cells accumulate RAD51 foci in response to DNA damage, a result of defective HR processing. These effects are reversed by ectopic expression of *MND1* or *PSMC3IP*, but not by a *PSMC3IP* mutant with an inability to form productive D-loops, which are critical for effective HR.

## Results

### Parallel CRISPR mutagenesis and interference screens identify *MND1* and *PSMC3IP* as highly penetrant determinants of PARPi sensitivity

To identify genetic determinants of PARPi sensitivity, we carried out parallel CRISPR mutagenesis (CRISPRn) and CRISPR interference (CRISPRi) chemosensitivity screens. In these screens, we used a PARPi resistant, HR proficient, non-tumor epithelial cell line with a previously engineered *TP53* mutation, MCF10A *TP53^-/-^*. We used *TP53* mutant MCF10A cells, as many cancer-associated mutations (such as *BRCA1* (Hakem et al., 1997; Hakem et al., 1996; Ludwig et al., 1997) often impair cellular fitness by invoking *TP53*-mediated cell cycle checkpoints and are thus better tolerated when *TP53* is inactivated. To facilitate CRISPR screening, we introduced into these cells either a doxycycline-inducible Cas9 transgene (Cas9) or a transgene expressing catalytically-inactive Cas9 (dCas9) fused to the KRAB transcriptional repressor (Gossen and Bujard, 1992). After confirming expression of either Cas9 or dCas9-KRAB in these cells (Figure 1A), we confirmed their PARPi resistance, finding that MCF10A *TP53^-/-^* cells were as resistant to PARPi as *BRCA1* mutant SUM149 triple negative breast tumor cell lines with a PARPi-resistance causing *BRCA1* reversion mutation (Bajrami et al., 2014; Drean et al., 2017) (Figure 1B, C). We tested this for two clinical PARPi, olaparib (Figure 1B) and talazoparib (Figure 1C).

**Figure 1.**
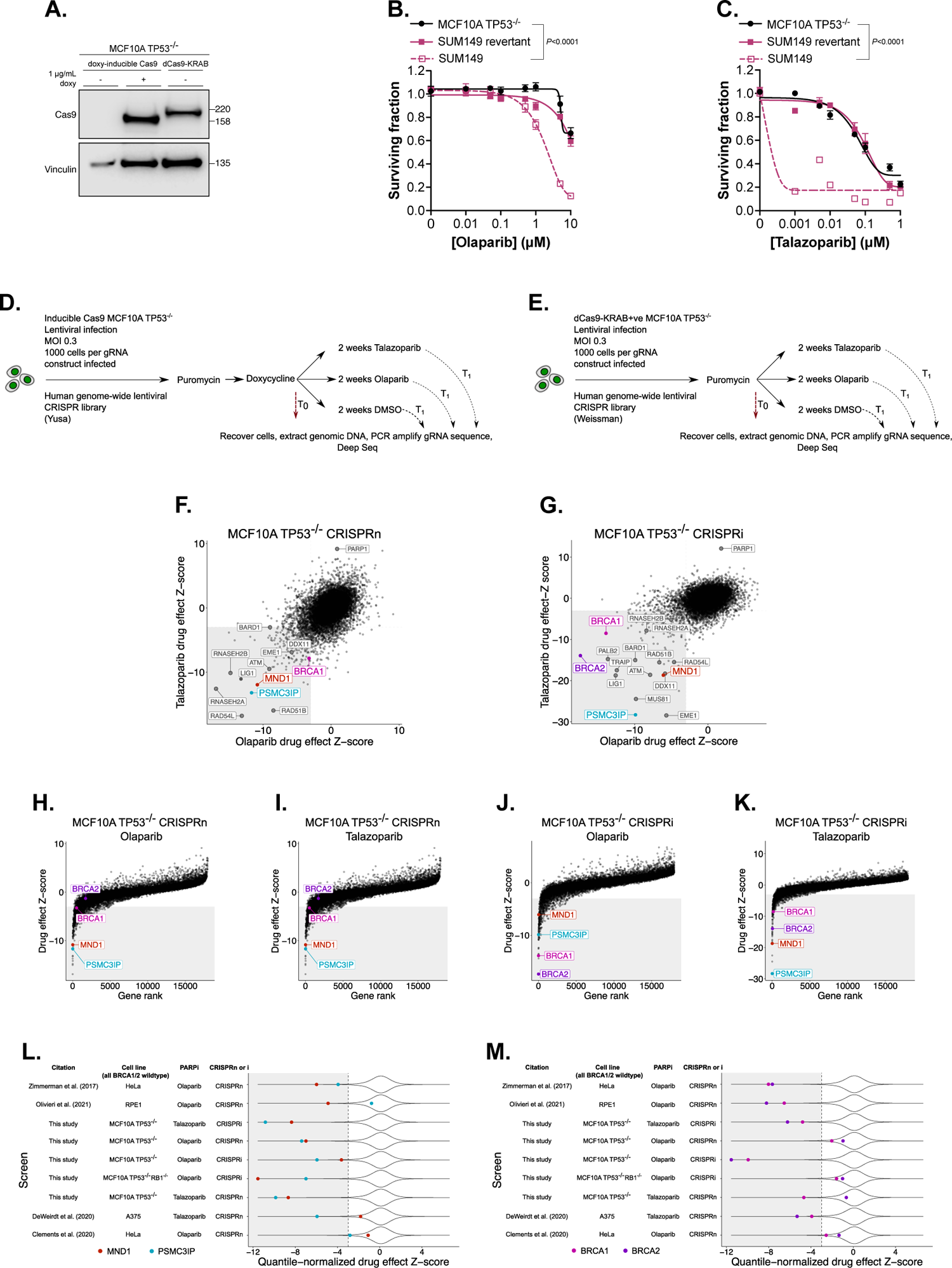
Parallel CRISPR mutagenesis and interference screens identify determinants of PARPi sensitivity. **A.** Western blot image of MCF10A *TP53^-/-^* cell lysates illustrating expression of either doxycycline-inducible Cas9 or catalytically-inactive Cas9 (dCas) fused to a KRAB transcriptional repressor (dCas9-KRAB). Vinculin was used as a loading control. Uncropped image shown in Supplementary Figure 12A. **B, C.** PARPi resistance in MCF10A *TP53^-/-^* cells. Dose/response survival curves are shown with surviving fractions at the indicated doses of olaparib (B) or talazoparib (C). Cells were plated in 384-well plates and exposed to PARPi for five continuous days, after which cell viability was quantified by CellTiter-Glo®. PARPi sensitive *BRCA1* mutant SUM149 and PARPi resistant *BRCA1* revertant SUM149 cells are shown as controls. Error bars represent SD from n=3 replicates. *P*-values were calculated via ANOVA with Tukey’s post-test. **D, E.** Schematics representing workflow for CRISPRn (D) and CRISPRi (E) screens. **F, G.** Data from genome-wide CRISPRn (F) and CRISPRi (G) screens. Scatter plots are shown with olaparib *vs.* talazoparib drug effect Z-scores. Genes with negative Z-scores represent PARPi sensitivity-causing effects (as shown by named DNA repair genes), whereas genes with positive Z-scores represent PARPi resistance-causing effects (e.g. *PARP1*)**. H-K.** Effects of *MND1* and *PSMC3IP* compared to effects elicited via CRISPRn or CRISPRi of *BRCA1* or *BRCA2*. Plots of genome-wide drug effect Z-scores compared to gene rank product based on MaGeCK and Z-score analysis (see Methods) from the screens described in Figure 1D, E. **L, M.** *MND1* and *PSMC3IP* are highly penetrant determinants of PARPi sensitivity. Violin plots of quantile normalized Z-score data (see methods) from nine different CRISPRn or CRISPRi screens for PARPi sensitivity, described either in this study or elsewhere (Clements et al., 2020; DeWeirdt et al., 2020; Olivieri and Durocher, 2021; Zimmermann et al., 2018). Data was analyzed using a consistent pipeline (see methods), to allow cross comparison. Quantile normalized Z-scores for *MND1* and *PSMC3IP* (L) or *BRCA1* and *BRCA2* (M) are highlighted. Each of the cell lines shown is a mitotic hTERT-positive/ALT-negative cell line.

For the CRISPRn screen, we mutagenized Cas9+ cells with a genome-wide single guide (sg)RNA library designed to target 18,010 protein coding genes (90,706 sgRNAs (Tzelepis et al., 2016) Figure 1D). In totality, 1 x 10^7^ cells were transduced at a multiplicity of infection of 0.3 (to ensure <1 sgRNA per cell), resulting in each sgRNA infecting at least 1000 cells, a representation that was maintained throughout the experiment. After removing non-transduced cells and a fraction of the cell population for later analysis (T_0_ sample), the resultant cell population was divided into three cohorts; these were either cultured in the presence of drug vehicle (DMSO), olaparib or talazoparib. Here, we used concentrations of PARPi sufficient to cause 20% reduction in the cell population (Surviving Fraction 80 concentration, SF_80_). Cells were continuously cultured in the presence of drug (or DMSO) for two weeks, at which point DNA from surviving cells was recovered (T_1_ sample). Using deep sequencing, we compared the relative enrichment or depletion of sgRNAs from T_0_ *vs.* T_1_ samples in both DMSO and PARPi exposed samples, and used these data to calculate normalized drug effect Z-scores (normZ (Colic et al., 2019)) for each gene; in this case CRISPR mutagenesis of genes with negative Z-scores caused enhanced PARPi sensitivity, with a Z-score threshold of <–3 being used to identify profound effects. The full set of Z-scores for the screens are provided in Supplementary Table 1, with the Z-scores for individual sgRNAs from CRISPRn and CRISPRi screens in Supplementary Tables 2 and 3, respectively. In our screens, including CRISPRi screens described below, we used an assessment of the depletion of sgRNA targeting core essential genes as a quality control measure (Hart et al., 2014), an approach which indicated that each of the screens was of sufficient quality to warrant further analysis (Supplementary Figure 1A-C, Supplementary Tables 2 and 3).

In parallel, we also conducted CRISPRi screens for olaparib or talazoparib sensitivity in dCas9-KRAB+ MCF10A *TP53^-/-^* cells (Figure 1E). These screens were carried out in a similar way to the CRISPRn screens, using an sgRNA library designed to silence 18,905 protein coding genes (104,535 sgRNAs (Horlbeck et al., 2016)). When we compared data from olaparib *vs.* talazoparib screens, we found both CRISPRi and CRISPRn screens to be highly reproducible (Figure 1F, G) providing confidence in their fidelity.

The cytotoxic effect of clinical PARPi, such as olaparib and talazoparib, is partially caused by trapping PARP1 on DNA (Krastev et al., 2021). Deletion of *PARP1* or *PARP1* mutations that prevent PARP1 trapping therefore cause PARPi resistance, both in *BRCA1* wild-type and *BRCA1* mutant cells (Pettitt et al., 2018). We noted that sgRNA designed to target *PARP1* gave one of the most profound PARPi resistance-causing effects in both CRISPRn and CRISPRi screens (Figure 1F, G). Previous focused shRNA screens (McCabe et al., 2006), genome-wide shRNA screens (Supplementary Table 4) (Bajrami et al., 2014) and genome-wide CRISPR-Cas9 screens for PARPi sensitivity (Clements et al., 2020; DeWeirdt et al., 2020; Jamal et al., 2022; Olivieri and Durocher, 2021) indicated that a number of different genes involved in HR enhance PARPi sensitivity when inactivated. This was also the case in our CRISPRn and CRISPRi screens. Some of the most profound PARPi sensitivity-causing effects were due to CRISPRi or CRISPRn targeting of the HR-associated genes *RAD51B, RAD54L, EME1, ATM, MUS81, PALB2, BRCA1, BARD1, BRCA2* and *DDX11* (Figure 1F, G) and when we carried out unbiased pathway annotation of “hits” in CRISPRn and CRISPRi screens, both olaparib and talazoparib screens identified HR as an enriched pathway (KEGG “homologous recombination” *p* values of 5.5 x 10^-10^, 1.1 x 10^-8^, 9.4 x 10^-16^, 1.9 x 10^-12^ for olaparib/CRISPRn, talazoparib/CRISPRn, olaparib/CRISPRi and talazoparib/CRISPRi screens respectively, Supplementary Table 5-8). In addition to the genes detailed above, additional genes involved in HR and double strand break repair also scored as “hits” in our screen (*ACTR5, ATM, ATRIP, AUNIP, CHAF1B, FAAP24 (C19orf40), FANCA, FANCD2, FANCE, FANCI, FANCL, FANCM, INO80, KIAA1524 (CIP2A), MCM8, MCM9, MRE11A, NBN, NSMCE1, NDNL2, PNKP, RAD50, RAD51, RAD51C, RAD51AP1, RBBP8 (CtIP), RMI1, RNF8, RNF168, SHFM1 (DSS1), SLX4, SMC4, SMC6, SFR1, STRA13 (CENPX), SLX4, SWI5, TELO, TONSL, TRAIP, USP1, WDR48, WRN, XRCC2, XRCC3*), recruitment and activity of the 9-1-1 complex (*ATAD5, RAD1, RAD9A, RAD17*), control of the DNA damage-induced S/G_2_ and G_2_/M checkpoints (*FOXM1*, *CCNB2*), chromatin remodeling complex components (*ACTL6A, BRD2, RBBP7, SMARCB1*), chromosome cohesion factors (*CHTF18, CHTF8, ESCO2, DSCC1*), base excision repair (*LIG1, LIG3, FEN1, UNG, APEX2, MUTYH*), nucleotide excision repair (*CUL4A, GTF2H2C, RFC4, LIG1, RPA3, POLD2, ERCC2, GTF2H3, RFC5, PCNA RFC1, CCNH, CETN2, GTF2H4, DDB1, POLE4, CDK7, ERCC3*), the PARP1 co-factor *C4orf27* (*HPF1*) and three *RNASEH2*-family genes known to control PARPi sensitivity by modulating levels of genomic uracil (*RNASEH2A, RNASEH2B, RNASEH2C* (Zimmermann et al., 2018)).

Our use of both CRISPRi and CRISPRn screens, and the use of two different clinical PARPi, allowed us to identify the most profound effects that were independent of the mode of gene perturbation or the PARPi used. As expected, this approach identified a number of HR-associated genes, but also identified two genes that encode a heterodimer classically involved in meiotic recombination, *MND1* (Meiotic Nuclear Division Protein 1 Homolog) and *PSMC3IP* (PSMC3 Interacting Protein, *HOP2*, Figure 1F, G, Supplementary Figure 1D-G). We also identified *MND1* and *PSMC3IP* depletion in a retroviral mutagenesis screen selecting HAP1 cells by ionizing radiation (IR) (Supplementary Figure 1H-K) (Francica et al., 2020), suggesting the effect of *MND1*-*PSMC3IP* inhibition was not specific to PARPi, but also caused sensitivity to other forms of DNA damage. When we examined each individual PARPi CRISPR-Cas9 screen, we found that the effect of targeting *MND1* or *PSMC3IP* on PARPi sensitivity was often comparable, or more profound, than that achieved by CRISPR-targeting of either *BRCA1* or *BRCA2* (Figure 1H-K, Supplementary Figure 1D-G). We also assessed the generality of these observations by re-analysis of published CRISPR screens carried out in different cell line backgrounds (HeLa, RPE1, A375) (Supplementary Table 9) and by carrying out an additional set of CRISPRn screens in another MCF10A derivative with an *RB1* tumor suppressor defect in addition to the *TP53* mutation (Supplementary Figure 1L-N, Supplementary Table 10). This analysis indicated that the relationship between CRISPR-targeting of *MND1* or *PSMC3IP* and PARPi sensitivity had comparable penetrance (if not more so) than the effect of targeting either *BRCA1* or *BRCA2* (Figure 1L, M), suggesting that in mitotic cells, the MND1/PSMC3IP heterodimer might be involved in the response to PARPi.

### MND1 and PSMC3IP control PARPi sensitivity in mitotic cells

MND1 and PSMC3IP proteins form a DNA binding heterodimer whose canonical function is associated with meiotic RAD51 or DMC1-mediated meiotic recombination (Chen et al., 2004; Tsubouchi and Roeder, 2002). As part of the meiotic recombination process, DNA double strand breaks (DSBs) are formed in the double helix by SPO11. These are then resected to generate exposed tracts of single stranded DNA (ssDNA), which are in turn bound by either DMC1 or RAD51, forming a helical presynaptic nucleoprotein filament. Using RAD51/DMC1 ATPase activity, the presynaptic filament invades duplex target DNA to form a heteroduplex DNA joint (D-loop), which is extended by DNA strand exchange and synthesis and then resolved to generate either crossover or non-crossover DNA recombinants (Crickard and Greene, 2018). As part of this meiotic process, the MND1/PSMC3IP heterodimer promotes meiotic interhomolog pairing by stabilizing the presynaptic filament and the capture of duplex DNA. Specifically, the N-terminal double-stranded DNA-binding functions of PSMC3IP/MND1 mediate synaptic complex assembly and the PSMC3IP C-terminus binds ssDNA and stabilizes the nucleoprotein filament (Zhao et al., 2014). PSMC3IP/MND1 also regulate ATP and DNA binding by RAD51 (Bugreev et al., 2014). Given the canonical role of MND1/PSMC3IP is in meiotic recombination, we were interested to understand why these genes might control response to DNA damaging agents, such as PARPi, in mitotic cells.

In addition to its role in meiotic recombination, there is some evidence that the MND1/PSMC3IP heterodimer also functions in mitotic cells, which predominantly carry out HR between sister chromatids as opposed to homologous chromosomes. MND1/PSMC3IP are expressed in tumor cell lines, particularly those that maintain telomeres via the alternative lengthening telomeres (ALT) pathway, a form of HR. As part of ALT, MND1/PSMC3IP promotes telomere clustering and RAD51-mediated recombination between otherwise geographically distant telomeres on different chromosomes (Cho et al., 2014; Dilley et al., 2016). To extend these observations, we analyzed gene expression and mass spectrometry proteomic data from human tumor cell lines (https://depmap.org) to assess the generality of MND1/PSMC3IP expression in mitotic cells. We found that in human tumor cell lines, MND1 and PSMC3IP mRNA and protein expression was relatively common (Supplementary Figure 2A, B) and not solely restricted to tumor cell lines that carry out telomere maintenance by ALT (Supplementary Figure 2C, D). For both MND1 and PSMC3IP, mRNA expression correlated with protein expression (Supplementary Figure 2E, F) and MND1 expression was highly correlated with PSMC3IP expression (Supplementary Figure 2D, G), consistent with the hypothesis that these two heterodimer components have a shared function in mitotic cells. Tumor expression of MND1 and PSMC3IP was also relatively common (Supplementary Figure 2H, I) and highly correlated, including in those tumor types where PARPi are used clinically (breast, serous ovarian, pancreatic adenocarcinoma and prostate adenocarcinoma, Supplementary Figure 2J-M).

On the basis of our CRISPR screen results, and the data indicating that MND1 and PSMC3IP are commonly expressed in mitotic tumor cells, we formally assessed whether *MND1* or *PSMC3IP* defects caused PARPi sensitivity. In CRISPRi experiments, we found that transduction of MCF10A *TP53^-/-^*cells expressing dCas9-KRAB with lentiviral constructs encoding sgRNA targeting *MND1* or *PSMC3IP* caused a reduction in MND1 or PSMC3IP mRNA levels and enhanced sensitivity to olaparib or talazoparib (Figure 2A-D, Supplementary Figure 3A, B), confirming the results observed in the screen. To further validate these results, we transfected MCF10A *TP53^-/-^*cells with Cas9-crRNA ribonucleoproteins targeting *MND1* or *PSMC3IP* and generated daughter clones with different *MND1* or *PSMC3IP* mutations (Figure 2E, F, Supplementary Figure 3C-J). *MND1* or *PSMC3IP* mutant clones were also sensitive to talazoparib (Figure 2G, H), a clinical PARPi known to effectively “trap” PARP1 on chromatin (Krastev et al., 2021; Murai et al., 2014). This was not the case for the poor PARP1-trapper, but effective PARP1 catalytic inhibitor, veliparib (Supplementary Figure 4A, B), suggesting that like PARPi *vs.* BRCA1/2 synthetic lethality (Shen et al., 2013), PARPi/MND1 or PARPi/PSMC3IP synthetic lethality might be more dependent upon PARP1 trapping than catalytic inhibition. Furthermore, *MND1* or *PSMC3IP* mutant clones were not sensitive to a small molecule ATR inhibitor (Supplementary Figure 4C, D) suggesting that the effect of PARPi did not necessarily extend to any agent that causes RF stress. PARPi sensitivity was not restricted to MCF10A *TP53^-/-^* cells, and was also seen in KB1P-G3B1 mouse mammary tumor cells (Barazas et al., 2019) grown *ex vivo* that were CRISPR-Cas9 mutagenized by *Mnd1* sgRNA (Figure 2I, J, Supplementary Figure 4E, F), which was partially rescued by Mnd1 overexpression (Figure 2K). PARPi sensitivity was also seen in *MND1* defective HAP1 cells (Figure 2L, M). We also confirmed sensitivity to IR in *MND1* or *PSMC3IP* mutant MCF10A *TP53^-/-^* cells (Supplementary Figure 4G, H) and in *Mnd1* defective KB1P-G3B1 cells (Figure 2N). Restoration of Mnd1 expression in *Mnd1* defective KB1P-G3B1 cells partially reversed radiosensitivity (Figure 2N, Supplementary Figure 4I). Taken together with our prior screen data, this suggested that the observed PARPi synthetic lethal effects (and also IR sensitivity) were relatively common effects in mitotic cells.

**Figure 2.**
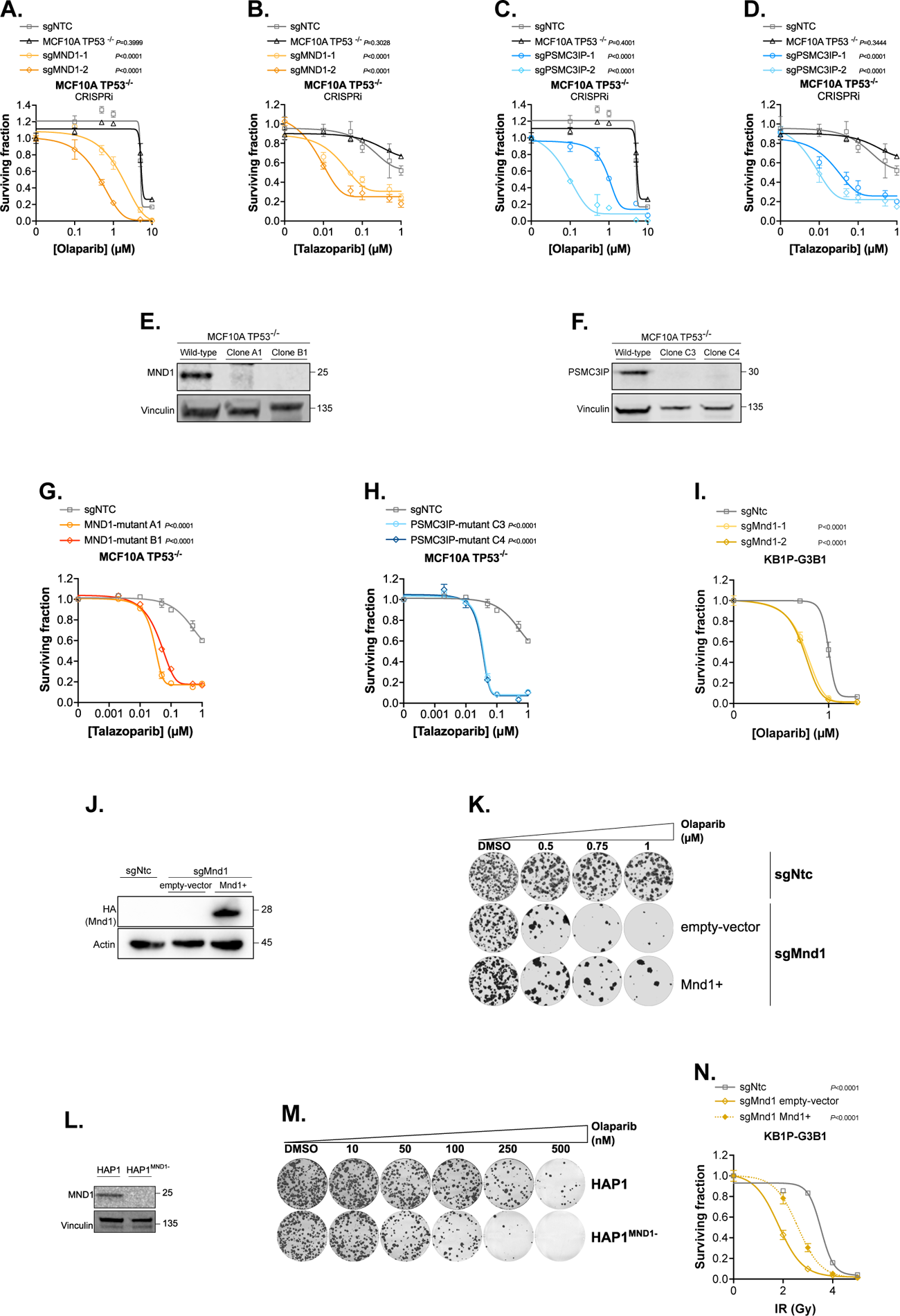
MND1 and PSMC3IP defects cause PARPi and ionizing radiation sensitivity in mitotic cells. **A-D.** Depletion of *MND1* or *PSMC3IP* using CRISPRi sensitized MCF10A *TP53^-/-^*cells to olaparib (A, C) and talazoparib (B, D). Dose/response survival curves are shown with surviving fractions at the indicated doses of PARPi. MCF10A *TP53^-/-^* cells expressing dCas9-KRAB were transduced with lentiviral constructs encoding sgRNA targeting *MND1* (sgMND1) (A, B) or *PSMC3IP* (sgPSMC3IP) (C, D). Cells were plated in 6-well plates (A, C) or 384-well plates (B, D) and exposed to PARPi for 14 continuous days (6-well plates) or five continuous days (384-well plates). Cell viability was quantified by CellTiter-Glo® and surviving fraction was calculated for each drug dose relative to DMSO-exposed cells. Error bars represent SD. *P*-values were calculated via ANOVA with Tukey’s post-test. **E**, **F.** Generation of MND1 or PSMC3IP mutant clones. MCF10A *TP53^-/-^* cells were transfected with non-targeting control or Cas9-crRNA ribonucleoproteins targeting *MND1* to generate daughter clones A1 and B1 (E) or *PSMC3IP* to generate daughter clones C3 and C4 (F). Western blot images demonstrating an almost complete absence of either MND1 (E) or PSMC3IP (F) in lysates extracted from mutant clones are shown. The antibodies used detect epitopes in the p.R82-E142 and p.P156-D216 regions of MND1 and PSMC3IP, respectively. The targeted epitopes are C-terminal to MND1 or PSMC3IP mutations generated. Vinculin was used as a loading control. Uncropped images shown in Supplementary Figure 12B, C. **G, H.** *MND1* (G) or *PSMC3IP* (H) mutant clones were more sensitive to talazoparib than wild type cells. Dose/response survival curves are shown with surviving fractions at the indicated doses of talazoparib. Cells were plated in 384-well plates and exposed to talazoparib for five continuous days, after which cell viability was quantified by CellTiter-Glo®. Error bars represent SD from n=3 replicates. *P*-values were calculated via ANOVA with Tukey’s post-test. **I.** *Mnd1* defective KB1P-G3B1 cells are more sensitive to olaparib than cells expressing non-targeting control. Dose/response survival curves are shown. KB1P-G3B1 cells were transduced with lentiviral constructs encoding sgRNA targeting *Mnd1* (either sgMnd1-1 or sgMnd1-2) or non-targeting control (sgNtc). Cells were plated in 6-well plates and exposed to olaparib for 11 continuous days, after which colonies were stained with crystal violet; colonies were quantified in an automated manner with macros using ImageJ. Representative image shown in Supplementary Figure 4E. Error bars represent SD. *P*-values were calculated via ANOVA with Tukey’s post-test. **J.** Western blot image of KB1P-G3B1 cell lysates illustrating restoration of Mnd1 expression via HA-tag in cells expressing a vector containing Mnd1 cDNA (Mnd1+), but not in cells expressing empty-vector. As indicated, KB1P-G3B1 cells express sgRNA targeting *Mnd1* (sgMnd1) or nontargeting control (sgNtc). Actin was used as a loading control. Uncropped image shown in Supplementary Figure 12D. **K.** Restoration of Mnd1 expression (Mnd1+) in *Mnd1* defective (sgMnd1) KB1P-G3B1 cells partially reversed PARPi sensitivity. Representative images of growth assays are shown. Cells were plated in 6-well plates and exposed to olaparib for 11 continuous days, after which colonies were stained with crystal violet. **L.** Western blot image demonstrating absence of MND1 in cell lysates extracted from *MND1*-defective, but not wild-type, HAP1 cells. Vinculin was used as a loading control. Uncropped image shown in Supplementary Figure 12E. **M.** HAP1 cells with defective *MND1* were more sensitive to olaparib than wild-type cells. Images of clonogenic assay are shown. Cells were plated in 6-well plates and exposed to olaparib for 14 continuous days, after which colonies were stained with sulforhodamine-B. **N.** *Mnd1* defective KB1P-G3B1 cells (sgMnd1 empty-vector) are more sensitive to IR compared to cells expressing non-targeting control (sgNtc). Restoration of Mnd1 in *Mnd1* defective KB1P-G3B1 cells (sgMnd1 Mnd1+) partially reversed radio-sensitivity. Dose/response survival curves are shown with surviving fractions at the indicated doses of IR. Cells were plated in 6-well plates and exposed to indicated dose of IR and then cultured for 11 continuous days, after which colonies were stained with crystal violet; colonies were quantified in an automated manner with macros using ImageJ. Representative image of colony formation is shown in Supplementary Figure 4I. Error bars represent SD. *P*-values were calculated via ANOVA with Tukey’s post-test.

### PARPi sensitivity in MND1/PSMC3IP defective cells is characterized by an increase in RAD51 foci and suppression of HR

The MND1-PSMC3IP heterodimer has been shown to facilitate meiotic RAD51 function in yeast (Tsubouchi and Roeder, 2002) and in cell-free *in vitro* assays, MND1-PSMC3IP catalyzes the binding of mouse and human RAD51 to nucleotides and DNA (Bugreev et al., 2014). The DNA lesions caused by PARP inhibitors and IR often activate HR and Ser-139 phosphorylation of histone variant H2AX (γH2AX), as well as the localization of the recombinase RAD51 to the site of DNA damage (Bryant et al., 2005; Farmer et al., 2005). Using proximity ligation assays (PLAs) in mitotic KB1P-G3B1 cells, we estimated the co-localization of Mnd1 with either Rad51 or γH2ax. We found that Mnd1 co-localized with Rad51 in the presence or absence of exogenous DNA damage, as previously seen in murine and human models (Chi et al., 2007; Petukhova et al., 2005; Pezza et al., 2006) (Figure 3A, Supplementary Figure 5A). Mnd1 co-localised with γH2ax solely upon exogenous DNA damage (Figure 3B, Supplementary Figure 5B). However, when we assessed the ability of RAD51 to localize to the site of DNA damage in *MND1* mutant MCF10A *TP53^-/-^* cells, we found that rather than seeing reduction in nuclear RAD51 foci (a phenotype normally associated with a HR defect, and radio- or PARPi-sensitivity (van Wijk et al., 2020), we observed significantly higher levels of RAD51 foci (Figure 3C, Supplementary Figure 5C, D). This was also true in KB1P-G3B1 mouse mammary tumor cells with an *Mnd1* defect (Supplementary Figure 6A, B), where ectopic expression of Mnd1 partially reversed this Rad51 foci increase. In *Mnd1* defective cells, Rad51 foci were also resolved with a delayed kinetic compared to controls (Figure 3D, Supplementary Figure 7A). We also observed a PARPi- or IR-induced increase of RAD51 foci in *PSMC3IP* mutant MCF10A *TP53^-/-^*cells (Figure 3E, Supplementary Figure 7B), consistent with the effects seen in *MND1* defective cells and is reminiscent of the persistence of nuclear RAD51 foci in *PSMC3IP* defective meiotic cells (Petukhova et al., 2003). We also saw a corresponding increase in γH2AX foci (Supplementary Figure 6C, D and 7B, C). To assess the impact of this increase in RAD51 foci on DNA repair by HR, we used a cell line with a synthetic HR reporter substrate (DR-GFP; (Gunn and Stark, 2012)) (Figure 3F) and found that either *MND1* or *PSMC3IP* gene silencing (Supplementary Figure 8A, B) caused a reduction in HR-mediated repair (Figure 3F, Supplementary Figure 8C). Taken together, the foci and DR-GFP data suggest that RAD51 function is defective in *MND1*/*PSMC3IP* deficient cells and they struggle to complete HR-mediated DNA repair.

**Figure 3.**
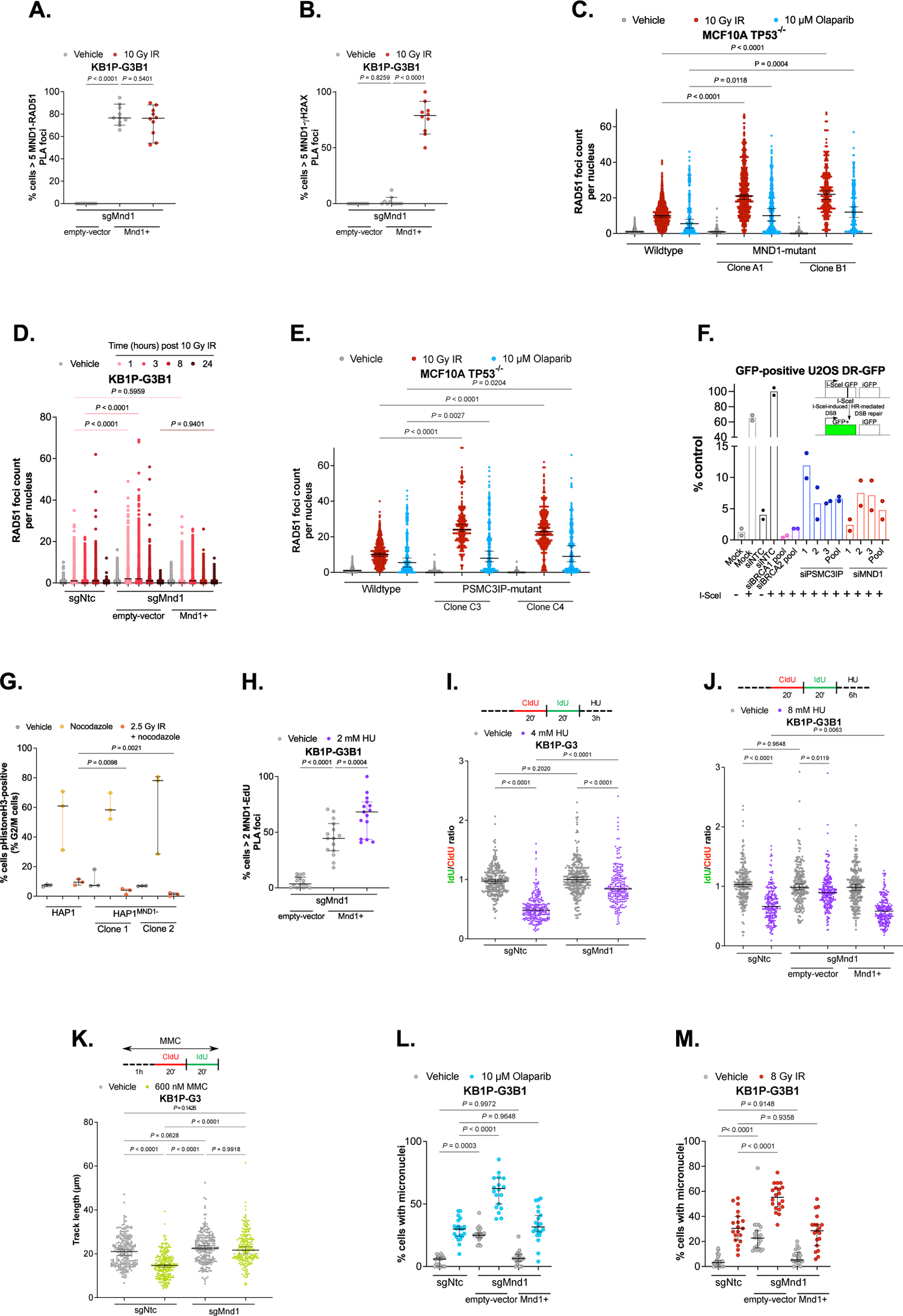
PARPi sensitivity in MND1/PSMC3IP defective cells is characterized by an increase in RAD51 foci and suppression of HR. **A, B.** Mnd1 co-localized with Rad51 in the presence or absence of exogenous DNA damage (A), but only co-localized with gH2ax upon exogenous DNA damage (B). Scatterplots are shown of percentage of cells with >5 PLA foci, normalized to the total number of cells imaged in multiple (n=10) fields of view. KB1P-G3B1 cells with a *Mnd1* defect (sgMnd1), either expressing empty-vector or a vector containing *Mnd1* cDNA (Mnd1+), were plated onto coverslips. Cells were exposed to 10 Gy IR or remained unexposed. Proximity ligation assays (PLAs) were performed following staining with anti-HA-tag (HA fused to Mnd1) and anti-RAD51 (A) or anti-gH2AX (B) antibodies. Error bars represent the median and 95% CI. *P*-values were calculated via ANOVA with Tukey’s post-test. Representative images are shown in Supplementary Figure 5A, B. **C.** Higher RAD51 foci levels were observed in *MND1* mutant cells compared to wild-type cells upon olaparib or IR exposure. Scatter plot of RAD51 foci count per nucleus (n=min. 157) in each indicated cell line is shown. MCF10A *TP53^-/-^*cells, either wild-type or with *MND1* defect (clones A1 and B1) were plated onto coverslips. Cells were either exposed to 10 µM olaparib and then fixed after 16 hours or 10 Gy IR and then fixed after 4 hours. Vehicle cells remained untreated and were fixed simultaneously with the olaparib- or IR-exposed samples. Cells were co-stained with anti-RAD51 and anti-gH2AX antibodies. gH2AX foci quantification in these samples in shown Supplementary Figure 5C. Error bars represent the median and 95% CI. *P*-values were calculated via ANOVA with Bonferroni’s post-test. **D.** Increased Rad51 foci levels and altered kinetics of Rad51 resolution were observed upon IR in *Mnd1* mutant cells compared to control cells. These phenotypes were partially reversed with ectopic Mnd1 expression. Scatter plot of RAD51 foci count per nucleus (n=min. 369) in each indicated cell line is shown. KB1P-G3B1 cells were plated onto coverslips, either expressing non-targeting control (sgNtc) or sgRNA targeting *Mnd1* (sgMnd1, expressing either empty-vector or vector containing *Mnd1* cDNA (Mnd1+). Cells were either exposed to 10 Gy IR and then fixed at the indicated timepoint or remained unexposed. Cells were stained with anti-RAD51 antibody. Error bars represent the median and 95% CI. *P*-values were calculated via ANOVA with Bonferroni’s post-test. Representative images shown in Supplementary figure 7A. **E.** Higher RAD51 foci levels were observed in *PSMC3IP* mutant cells compared to wild-type cells upon PARPi or IR exposure. Scatter plot of gH2AX foci count per nucleus (n= min. 181) in each indicated cell line is shown. MCF10A *TP53^-/-^* cells, either wild-type or with *PSMC3IP* defect (clones C3 and C4) were plated onto coverslips. Cells were either exposed to 10 µM olaparib and then fixed after 16 hours or 10 Gy IR and then fixed after 4 hours. Vehicle cells remained untreated and were fixed simultaneously with the olaparib-or IR-exposed samples. Cells were co-stained with anti-RAD51 and anti-gH2AX antibodies. gH2AX foci quantification in these samples is shown in Supplementary Figure 7C. Error bars represent the median and 95% CI. *P*-values were calculated via ANOVA with Bonferroni’s post-test. Representative images shown in Supplementary Figure 7B. **F.** *MND1* or *PSMC3IP* silencing reduced HR-mediated repair. Bar plot of % GFP+ cells relative to cells transfected with both non-targeting control siRNA (siNTC) and I-*Sce*I is shown. Schematic of assay shown in right panel. U2OS DR-GFP cells (Gunn and Stark, 2012) were transfected with siRNAs targeting *MND1, PSMC3IP* or non-targeting control, prior to expression of I-*Sce*I. A proportion of cells remained untransfected (mock) and another proportion of cells transfected with indicated siRNAs were not transfected with I-*Sce*I for controls of background GFP positivity. siRNA-mediated silencing of *BRCA1* or *BRCA2* were used as positive controls for HR deficiency. GFP+ cells were analyzed by flow cytometry. Representative FACS scatterplots shown in Supplementary Figure 8C. **G.** MND1 is required for cell cycle progression following DNA damage. Scatter plot of % phospho(p)HistoneH3+ cells is shown. pHistoneH3 was used as a mitotic marker (Wei et al., 1998). Cells were either left untreated, exposed to 250 ng/mL nocodazole for 16 hours, or exposed to 2.5 Gy IR 30 minutes prior to the 16-hour nocodazole exposure. Cells were fixed and stained with propidium iodide (PI) and anti-pHistoneH3. Representative FACS scatterplots shown in Supplementary Figure 9D. **H.** Mnd1 co-localizes with EdU-labelled nascent DNA, which is further increased by hydroxy urea (HU)-induced RF stalling. Scatterplots of SIRF assay are shown of cells with >2 PLA foci, normalized to the total number of cells imaged in multiple (n=15) fields of view. KB1P-G3B1 cells with *Mnd1* defect, either expressing empty-vector or vector containing Mnd1 cDNA (Mnd1+), were plated on coverslips with EdU. Cells were either exposed to 2 mM hydroxy urea (HU) for two hours or remained unexposed. PLAs were performed following staining with anti-HA-tag (tagged to Mnd1) and anti-biotin antibodies. Error bars represent the median and 95% CI. *P*-values were calculated via ANOVA with Tukey’s post-test. Representative images shown in Supplementary Figure 9G. **I, J.** Mnd1 is important for Brca1-independent RF degradation upon replication-blocking DNA damage. Schematic of DNA fiber assay performed in *Brca1*-deficient KB1P-G3 cells (I) and *Brca1*-proficient KB1P-G3B1 cells (J), as described in (Schmid et al., 2018) with a few modifications (detailed in upper panels). Scatter plots showing quantification of IdU/CldU ratio of at least n=120 fibers per sample (lower panels). Pulse-labelling followed by RF stalling via hydroxy urea (HU) resulted in an increased track length ratio of *Mnd1*-mutant cells (sgMnd1) compared to non-targeting control (sgNtc). Track length ratio was restored to wild-type levels with reconstitution of *Mnd1* cDNA (Mnd1+) (J). Error bars represent the median and 95% CI. *P*-values were calculated via ANOVA with Tukey’s post-test. **K.** Mnd1 is important for RF slowing, specifically fork reversal, upon replication-blocking DNA damage. Schematic of DNA fiber assay performed in Brca1-deficient KB1P-G3 cells, as described in (Schmid et al., 2018) with a few modifications (detailed in upper panel). Scatter plot showing quantification of total track lengths of at least n=120 fibers per sample (lower panel). Pulse-labelling followed by RF stalling via MMC resulted in increased RF progression, evidenced by increased track length, of *Mnd1*-mutant (sgMnd1) cells compared to non-targeting control (sgNtc). Error bars represent the median and 95% CI. *P*-values were calculated via ANOVA with Tukey’s post-test. **L, M.** *Mnd1* loss increases micronuclei formation upon exposure to olaparib (L) or IR (M), which is reversed with ectopic Mnd1 expression. Scatterplots of % cells with micronuclei in each indicated sample are shown (n=20). KB1P-G3B1 expressing either sgRNA targeting *Mnd1* (sgMnd1) or non-targeting control (sgNtc) were plated onto coverslips. *Mnd1*-deficient cells either express an empty-vector or a vector containing *Mnd1* cDNA (Mnd1+). Cells were either exposed to 8 Gy IR, 10 µM olaparib for 16 hours or remained untreated. Error bars represent the median and 95% CI. *P*-values were calculated via ANOVA with Tukey’s post-test. Representative images shown in Supplementary Figure 10O. Scatterplots showing *Psmc3ip* loss also increases micronuclei formation upon exposure to olaparib or IR shown in Supplementary Figure 10K, L) with representative image in Supplementary Figure 10R.

In response to DNA damage, RAD51 has been shown to contribute to sister chromatid exchange (SCE) (Lambert and Lopez, 2001), a crossover event that resolves Holliday junctions that shares some similarities with meiotic recombination (Lingg et al., 2022). To investigate SCE, we used HAP1 cells in which a clear increase in SCE is visible following olaparib– or IR-induced DNA damage (Supplementary Figure 9A-C). *MND1* knockout did not alter the rates of SCEs in untreated, olaparib- or IR-treated cells (Supplementary Figure 9B, C). While studying SCE in the *MND1* knockout cells, we observed fewer metaphase spreads in IR-treated cells, suggesting that these cells do not efficiently enter mitosis. We therefore measured mitotic entry after IR-induced G_2_ arrest. In contrast to HAP1 *MND1* wild-type cells, only few *MND1*-knockout HAP1 cells entered mitosis and most cells remained stuck in the G_2_ phase of the cell cycle (Figure 3G, Supplementary Figure 9D). In addition, a greater proportion of *Mnd1* defective KP1P-G3B1 cells were in G2/M compared to sgNtc even in untreated conditions, a phenotype which was rescued by ectopic expression of *Mnd1* (Supplementary Figure 9E, F). These data suggest that *MND1* is required for cell cycle progression, an effect that is enhanced following DNA damage. This may be explained by an altered response of the progressing RF to DNA damage. In this context, RAD51 has been identified to mediate RF reversal in a BRCA1/2-independent fashion, a mechanism which processes stalled RFs and appears to protect cells against genotoxic stress (Mijic et al., 2017; Zellweger et al., 2015). Moreover, in a BRCA1/2-dependent manner, RAD51 filament formation is required for its protective effect on the regressed arm, allowing PARP1/RECQ1-regulated restart of reversed RFs (Mijic *et al*., 2017; Schlacher et al., 2012; Zellweger et al., 2015). High concentrations of PARPi accelerate RF progression (Maya-Mendoza et al., 2018). Based on these findings, we hypothesized that the MND1-PSMC3IP heterodimer contributes to RAD51 function at RFs. This prompted us to first test whether MND1 is present at RF using the *in situ* analysis of protein interactions at DNA RF (SIRF) assay (Roy and Schlacher, 2019). Indeed, we found Mnd1 to co-localize with EdU-labelled nascent DNA in KB1P-G3B1 cells, an interaction further increased by hydroxy urea (HU)-induced RF stalling (Figure 3H, Supplementary Figure 9G). To investigate whether defective *Mnd1* affects the stability of stalled RFs, we first used *Brca1*- and *Tp53*-deficient KB1P-G3 cells (Barazas et al., 2019). As expected, pulse-labelling with CldU and IdU followed by RF stalling using 4 mM HU resulted in a significant reduction in the IdU/CldU track length ratio, indicating nucleolytic degradation of the nascent DNA of reversed RFs (Figure 3I). This was consistent with previous findings that BRCA1 stabilizes stalled forks (Schlacher *et al*., 2012). Interestingly, this fork degradation phenotype was reversed in *Mnd1*-mutant cells (Figure 3I). We then investigated the effect of Mnd1 on RF stability in isogenic *Brca1* proficient KB1P-G3B1 cells. Due to the presence of Brca1, a high concentration of 8 mM HU was needed to generate the RF intermediates that eventually become degraded (Figure 3J). In these cells, RF degradation was rescued by loss of Mnd1; an effect that was reversed by reconstitution of the *Mnd1* cDNA (Figure 3J). These data suggested that the effect of MND1 on RF stalling was BRCA1-independent. A reason for the lack of HU-mediated degradation in *MND1* defective cells may be a defect in RF reversal, a BRCA1/2-independent effect previously described in RAD51-deficient cells (Mijic et al., 2017; Qiu et al., 2021). Potent RAD51-dependent slowing of RF and their reversal is achieved by MMC treatment (Zellweger et al., 2015). Indeed, when we exposed KB1P-G3 cells to 600 nM MMC for 2 hours, we observed a clear slowing of RF progression (Figure 3K). Interestingly, *Mnd1* loss counteracted this fork slowing, consistent with a defect in RF reversal (Figure 3K). In fact, RF progression in *Mnd1*-mutant cells was slightly higher than in the non-targeting control cells, even in the absence of drug (vehicle) (Figure 3K). These data suggested that MND1 is important for RF slowing upon replication-blocking DNA damage. In its absence, unrestrained RF progression may result in accumulation of toxic DNA damage. Moreover, as *Mnd1* deficient cells appear to get stuck in the G_2_ phase of the cell cycle (presumably to deal with persistent DNA damage), we hypothesize that the MND1-PSMC3IP heterodimer may have its major protective role in RAD51-mediated HR that does not result in cross-over events.

We then investigated whether *MND1* loss is epistatic with defects in BRCA1 and BRCA2, two key proteins involved in HR. For this purpose, we exposed *Mnd1*-mutated *Brca1*- and *Tp53*-deficient KB1P-G3 cells to olaparib or IR (Supplementary Figure 10A-E). Compared to control cells and our prior observations in *Brca1/Brca2* wild-type cells (Figure 2I), we did not observe further PARPi or IR sensitization when *Mnd1* was CRISPR-Cas9 targeted. *Mnd1* mutation in the *Brca2*-*Tp53*-deficient mouse mammary tumor cell line KB2P3.4 (Evers et al., 2008) also did not elicit further PARPi sensitivity (Supplementary Figure 10F-J). This epistatic relationship was consistent with a crucial role for MND1 in HR in mitotic cells. Indeed, we found an increase in micronuclei formation in *Mnd1*- or *Psmc3ip*-deficient KB1P-G3B1 cells exposed to olaparib or IR (Figure 3L, M, Supplementary Figure 10K, L). These micronuclei represent broken chromosome parts and not mis-segregation of whole chromosomes, as they were negative for the centromere marker CENP-B (Supplementary Figure 10M-R). Since RAD51 nucleofilament formation is rather downstream in the HR pathway, we hypothesized that *MND1* depletion should also sensitize *BRCA1*- and *TP53*-deficient cells that acquired PARPi resistance by loss of *TP53BP1*, which restores HR. Indeed, when we mutated *Mnd1* in *Brca1^-/-^ Tp53bp1^-/-^* cells, these regained olaparib sensitivity and showed increased levels of RAD51 foci upon IR or PARPi exposure (Supplementary Figure 11).

Together, these data strongly suggest that the main control of PARPi response of the MND1-PSMC3IP heterodimer in mitotic cells is due to its role in supporting RAD51-mediated D-loop formation.

### PARPi sensitivity is reversed by wild-type PSMC3IP but not a p.Glu201del mutant associated with female gonadal dysgenesis and a D-loop defect

To functionally confirm the relevance of a D-loop defect in PARPi response, we made use of a previously described p.Glu201del mutant of PSMC3IP (Figure 4A). Premature truncating mutations or a deletion of p.Glu201 in PSMC3IP have been associated with XX female gonadal dysgenesis (XX-GD), a condition characterized by the lack of spontaneous pubertal development, primary amenorrhea, uterine hypoplasia, and hypergonadotropic hypogonadism (Peng et al., 2013). Although the p.Glu201del mutation (in the C-terminus of PSMC3IP) does not diminish the interaction of the MND1/PSMC3IP heterodimer with DNA, the interaction with RAD51 is impaired as is the ability to promote D loop formation (Zhao and Sung, 2015). We found that re-expression of wild-type *PSMC3IP* reversed PARPi sensitivity in *PSMC3IP*-depleted cells, establishing causality of *PSMC3IP* in PARPi response (Figure 4A, B). Interestingly, expression of a p.Glu201del mutant version of *PSMC3IP* did not demonstrate this phenotype, but rather further sensitized the cells to PARPi (Figure 4A, B). Expression of the *PSMC3IP* p.Glu201del mutant also sensitized MCF10A *TP53^-/-^* cells with wild-type *PSMC3IP* to PARPi (Figure 4A, B), consistent with this mutation acting as a dominant negative (Zhao and Sung, 2015). Consistent with our aforementioned observation that PARPi sensitivity in *PSMC3IP*-mutant cells is associated with an increase in RAD51 foci, expression of *PSMC3IP* p.Glu201del resulted in increased RAD51 foci formation upon IR or PARPi treatment (Figure 4C-G). This was true for *PSMC3IP*-depleted cells as well as cells with normal levels of *PSMC3IP*. As expected, re-expression of wild-type *PSMC3IP* reversed RAD51 foci formation in *PSMC3IP*-depleted cells, establishing causality of *PSMC3IP* in RAD51 foci formation (Figure 4C-G). We hypothesized that the increased RAD51 nucleoprotein formation in *MND1* and *PSMC3IP* mutant cells exposed to PARPi might be the key cytotoxic event. Consistent with this, inhibition of RAD51 with the previously-described RAD51 inhibitor B02, which inhibits the single- and double-stranded DNA binding and strand-exchange activity of RAD51 (Huang et al., 2012; Huang et al., 2011), partially reversed the PARPi sensitivity phenotype in both *MND1* and *PSMC3IP* mutant cells (Figure 4H,I).

**Figure 4.**
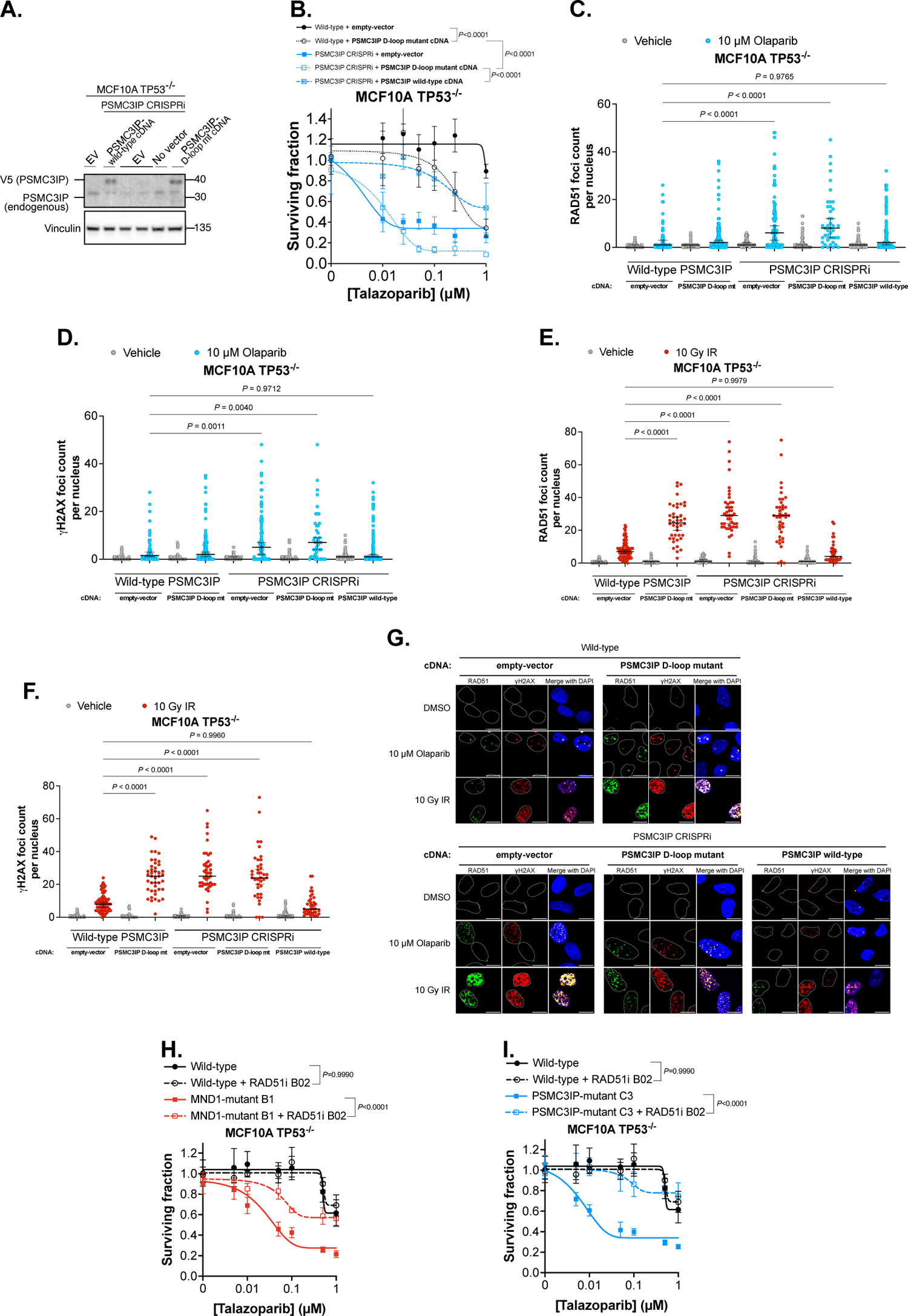
PARPi sensitivity is reversed by wild-type PSMC3IP but not a D-loop defective p.Glu201del mutant associated with female gonadal dysgenesis. **A.** Western blot image of MCF10A *TP53^-/-^* dCas9-KRAB cells (with or without sgRNA targeting *PSMC3IP*) with ectopic expression of either wild type or p.Glu201del (D-loop mutant) PSMC3IP. Vinculin was used as a loading control. Uncropped image shown in Supplementary Figure 12F. **B.** The PARPi-sensitivity phenotype associated with *PSMC3IP* defect is reversed with wild-type *PSMC3IP*, but not *PSMC3IP* p.Glu201del mutant. Dose/response survival curves are shown with surviving fractions at the indicated doses of talazoparib. Wild-type MCF10A *TP53^-/-^* cells expressing *PSMC3IP* p.Glu201del mutant are also more sensitive to PARPi, compared to cells expressing empty-vector. MCF10A *TP53^-/-^* dCas9-KRAB cells expressing non-targeting control (wild-type) or sgRNA targeting *PSMC3IP* (PSMC3IP CRISPRi) were transduced with lentiviral constructs encoding expression vector containing *PSMC3IP* cDNA, either wild-type *PSMC3IP* (*PSMC3IP* wild-type cDNA) or *PSMC3IP* p.Glu201del (PSMC3IP D-loop mutant cDNA). Wild-type or PSMC3IP CRISPRi cells expressing empty-vector were used as controls. Cells were plated in 96-well plates and exposed to talazoparib for ten continuous days. Cell viability was quantified by CellTiter-Glo® and surviving fraction was calculated for each drug dose relative to DMSO-exposed cells. Error bars represent SD from n=3 replicates. *P*-values were calculated via ANOVA with Tukey’s post-test. **C-G.** Higher RAD51 (C, E) and gH2AX (D, F) foci levels were observed in PSMC3IP CRISPRi cells upon olaparib (C, D) or IR (E, F) exposure, which was partially reversed with expression of wild-type *PSMC3IP*, but not *PSMC3IP* p.Glu201del D-loop mutant. Scatter plot of RAD51 foci count per nucleus (n= min. 41) in each indicated cell line is shown in C-F. Cells were plated onto coverslips and either exposed to 10 µM olaparib and then fixed after 16 hours or 10 Gy IR and then fixed after 4 hours or remained untreated. Cells were co-stained with anti-RAD51 and anti-gH2AX antibodies. Error bars represent the median and 95% CI. *P*-values were calculated via ANOVA with Bonferroni’s post-test. Representative images are shown in G; scale bar = 10 µm. **H, I.** RAD51 inhibition reverses the PARPi sensitivity phenotype in both *MND1* (H) and *PSMC3IP* (I) mutant cells. Dose/response survival curves are shown with surviving fractions at the indicated doses of talazoparib. Cells were plated in 384-well plates and exposed to 25 µM small molecule RAD51 inhibitor B02 for one hour prior to talazoparib addition. Cells were exposed to PARPi for five continuous days. Cell viability was quantified by CellTiter-Glo® and surviving fraction was calculated for each drug dose relative to DMSO-exposed cells. Error bars represent SD from n=3 replicates. *P*-values were calculated via ANOVA with Tukey’s post-test.

## Discussion

In this study, we identified *PSMC3IP* and *MND1* as key determinants of PARPi and IR response in mitotic cells. Our data are consistent with a role for the PSMC3IP-MND1 heterodimer in controlling RAD51 nucleofilament-mediated D-loop formation in mitotic cells, in addition to their role in inter-chromosomal recombination in meiotic cells. Some evidence that the MND1-PSMC3IP heterodimer also functions in mitotic cells came from the observation that these proteins contribute to ALT (Cho et al., 2014; Dilley et al., 2016): the MND1-PSMC3IP heterodimer promotes telomere clustering and RAD51-mediated recombination between distant telomeres. Our data show that both MND1 and PSMC3IP are also expressed in mitotic cells which do not use ALT. Hence, the MND1-PSMC3IP heterodimer seems to have a more general supportive role of RAD51-associated functions in mitotic cells that goes beyond ALT.

We found that *PSMC3IP*- or *MND1*-deficient cells have an increased PARPi sensitivity that is comparable to the loss of *BRCA1* or *BRCA2*. The knockout of *Mnd1* in *Brca1*- or *Brca2*-deficient cells did not further increase PARPi sensitivity, consistent with an epistatic role in the HR pathway. An important difference to *BRCA1/2* defects is that cells lacking *PSMC3IP* or *MND1* function have persisting RAD51 foci formation following DNA damage. In contrast, *BRCA1/2*-mutant cells do not form RAD51 foci and the lack of RAD51 foci formation is used as a surrogate marker for HR deficiency and to predict PARPi response (Cruz et al., 2018; Llop-Guevara et al., 2021; van Wijk et al., 2020). Here, we present an example of prolonged RAD51 foci formation, which is also an outcome of dysfunctional HR. This suggests that as well as assessing the loss of RAD51 foci in order to predict PARPi sensitivity, assessing the kinetics of RAD51 foci formation and resolution might also be important, as prolonged RAD51 foci formation may also indicate HR defects. It would be interesting to investigate whether this phenotype can be observed in tumors of patients that respond to PARPi, despite the presence of RAD51 foci.

Another feature that we observed in *MND1*-deficient cells is a defect in RF slowing following DNA damage. This is consistent with a contribution of MND1 to RF reversal, another RAD51-associated function (Mijic et al., 2017; Qiu et al., 2021). In the absence of RF slowing following DNA damage, unrestrained RF progression may increase the amount of DNA damage, including DSBs. The HR-mediated repair of DSB is clearly a major function that both *MND1* and *PSMC3IP* have in mitotic cells, as shown with the DR-GFP reporter assay. Within the HR pathway, we conclude that an impaired D-loop formation is responsible for the HR defect and toxic RAD51 foci formation in *MND1* or *PSMC3IP* defective cells. This is based on our experiments using the p.Glu201del mutant of *PSMC3IP*, a mutation that does not alter the interaction of the MND1-PSMC3IP heterodimer with DNA but which does impair the interaction with RAD51 and its ability to promote D loop formation (Zhao and Sung, 2015). In contrast to wild-type *PSMC3IP*, the p.Glu201del mutant does not recue PARPi-induced prolonged RAD51 foci formation and PARPi sensitivity. These conclusions are strengthened with our experiments demonstrating rescue of PARPi sensitivity of *PSMC3IP*-defective cells using small molecule RAD51 inhibitor, B02, which specifically inhibits single- and double-stranded DNA binding and strand-exchange activity of RAD51. PARPi sensitivity of *MND1-*defective cells was also rescued with B02. Interestingly, this RAD51 inhibitor has been previously shown to increase PARPi sensitivity of HR proficient TNBC cell lines, but not HR-proficient non-TNBC cell lines, such as MCF10A which is the main breast model used in this study (Shkundina et al., 2021).

Overall, our data contributes to the mechanistic insight to our understanding of how PARPi response is controlled and how MND1-PSMC3IP regulate the response to DNA damage in mitotic cells, outside of their role in ALT.

## Materials and Methods

### Cell Lines

MCF10A *TP53^-/-^* cells and MCF10A *TP53^-/-^ RB1^-/-^* daughter cells generated by CRISPR-Cas9 mutagenesis were purchased from Horizon. Cells were cultured in DMEM/Ham’s F-12 according to manufacturer’s instructions. DR-GFP U2OS (kindly gifted by Jeremy Stark (City of Hope, USA)), HEK293T (ATCC), CAL51 (DSMZ) and MDAMB-231 (ATCC) were maintained in Dulbecco’s Modified Eagle Medium (DMEM, Gibco) supplemented with 10% FBS. SUM149 cells (Asterand Bioscience) were maintained in Ham’s F-12 medium supplemented with 5% FBS, 10 µg/mL insulin and 1 µg/mL hydrocortisone. The KB1P-G3 cell line was previously established from a *K14cre;Brca1^F/F^;Trp53^F/F^*(KB1P) mouse mammary tumor and cultured as described by (Jaspers et al., 2013). The KB1P-G3B1 cell line was derived from the KB1P-G3 cell line which was reconstituted with human *BRCA1* by (Barazas et al., 2019). The *Tp53bp1* knock out KB1P-G3 line was generated by (Barazas et al., 2019). The KB2P-3.4 cell line was previously established from a *K14cre;Brca2^F/F^;Trp53^F/F^* (KB2P) mouse mammary tumor as described by (Evers et al., 2008). KB-derived cell lines were grown in Dulbecco’s Modified Eagle Medium/Nutrient Mixture F-12 supplemented with 10% FCS and 5 µg/mL Insulin 5 ng/mL cholera toxin and 5 ng/mL murine epidermal growth-factor (EGF, Sigma, #E4127). The HEK293FT cell line (RRID: CVCL_6911), as well as the Phoenix-ECO cell line (RRID:CVCL_H717), were cultured in DMEM (Gibco) supplemented with 10% FCS. HAP1 cells for SCE experiments were a kind gift from Thijn Brummelkamp, NKI. HAP1 cells used to generate olaparib dose/response survival curves were purchased from Horizon. HAP1 cells were cultured in IMDM containing 10% fetal bovine serum, 1% penicillin-streptomycin, and 1 mM L-glutamine (all reagents from Gibco). Tissue culture was carried out under standard conditions (37°C, 5% CO_2_), except for KB1P-G3 and KB2P3.4 cells which were cultured under low oxygen conditions (3% O_2_). Testing for mycoplasma contamination was performed on a regular basis.

MCF10A *MND1* and *PSMC3IP* mutant cell lines were engineered using the Edit-R Gene Engineering System (GE Dharmacon). Cells were seeded at a density of 1×10^6^ cells/well in 6-well plates. After 24 hours, cells were transfected with 40 µM Edit-R Cas9 nuclease protein NLS (CAS11729) mixed with 20 µM 2X crRNA and 10 µM tracrRNA using Lipofectamine CRISPRMAX transfection reagent, according to manufacturer’s instructions (Life Technologies). Target sequences for crRNA used: GCTGACCTTCAAGTCCTAGA and GTGAGGTTGAACACTTACTT to target *PSMC3IP*, CTTGCATGAAGAGCTTTACT and CGGAACTTCTAATTATTATT for *MND1* targeting, GATACGTCGGTACCGGACCG for non-targeting control. Four days after transfection, cells were FACS-sorted into 96-well plates at one cell per well in drug-free medium. Targeted genome modifications were analysed by Sanger sequencing of PCR products cloned into pCR-TOPO-blunt (Life Technologies). Constructs were introduced into MCF10A TP53*^-/-^* cells expressing inducible Cas9, which was generated by lentiviral transduction with hEF1a-Cas9 (#CAS11229, Dharmacon). Cas9 expression was induced with 1 µg/mL doxycycline.

MCF10A *MND1* and *PSMC3IP* CRISPRi cell lines were generated by cloning sgRNAs into the BbsI site of the pKLV5-U6sgRNA5-PGKPuroBFP (Addgene # 50946), as previously described (Tzelepis et al., 2016). sgRNA sequences are as follows: sgMND1-1: GCGGCGAAGCCCACACACTA; sgMND1-2: GGTAGCCTCAGTCCTTACCA; PSMC3IP-1: GCGGGAAAGGCGATGAGTAA; PSMC3IP-2: GAAGCTGCGGCGGGAGGTAA. These constructs were introduced into cells generated by lentiviral transduction of MCF10A TP53-/- cells with lenti-BLAST-dCas9-KRAB (Addgene, #89567) followed by selection with 10 μg/mL blasticidin.

In order to generate cells expressing a *PSMC3IP* mutant associated with D-loop defect (and XX-GD), a human PSMC3IP ORF (Dharmacon) was PCR-amplified using primers designed to result in a deletion of glutamic acid at amino acid positive 201 Fw-GCAAGAAGCAGTTCTTTGAGGTTGGGATAGAGACGGATGAAG Rev-CTCAAAGAACTGCTTCTTGCTCTTG. In-fusion reaction was performed to re-circularise the vector. *PSMC3IP* p.Glu201del or wild-type *PSMC3IP* cDNA was cloned into pLX302 (Addgene #25896) expression vector. These constructs were introduced into wild-type MCF10A *TP53^-/-^*cells or MCF10A *TP53^-/-^ PSMC3IP* CRISPRi cell lines (generated as per aforementioned procedure).

Generation of CRISPR/SpCas9 plasmids targeting KB1P-G3B1, KB1P-G3 and KB2P-3.4 cell lines, were performed using a modified version of the lentiCRISPR v2 backbone (Addgene #52961), in which a puromycin resistance ORF was cloned under the hPGK promoter. sgRNA sequences are as follows for KB1P-G3, KB1P-G3B1, KB2P-3.4 cell lines. Non-targeting control: TGATTGGGGGTCGTTCGCCA; sgMnd1-1: GACAAACATACCGTCTCTTGC; sgMnd1-2: GTCATGCCAGGAAGCGCAAGT. sgPscm3ip: GTAGGTTTCCGAACACGTCCT sgRNA sequence was introduced into KB1P-G3B1 for *Psmc3ip* targeting. The target sites modifications of the polyclonal cell pools were analyzed by TIDE analysis as follows; extracted genomic DNA was PCR-amplified and submitted with corresponding forward primers for Sanger sequencing to confirm target modifications using the TIDE algorithm (Brinkman et al., 2014).

For the HAP1 cell line used for SCE experiments, pDG459 backbone (Addgene #100901) carrying two sgRNAs was generated. Sequences are as follows for HAP1 cell lines. Non-targeting control: TGATTGGGGGTCGTTCGCCA; sgMnd1-1: GAGAAAAGAGAACTCGCATGA; sgMnd1-2: AAGCTTAGTTGATGATGGTA. All construct sequences were verified by Sanger sequencing.

### CRISPR screens

Inducible Cas9 MCF10A *TP53^-/-^* cells were generated for the CRISPRn screen by lentiviral transduction of MCF10A *TP53^-/-^*cells with hEF1a-Cas9 (#CAS11229, Dharmacon). Followed selection with 10 μg/mL blasticidin, Cas9-expressing cells were infected at multiplicity of infection (MOI) 0.3, with a previously published genome-wide human lentiviral CRISPR library (Tzelepis et al., 2016). The library contains 90,709 sgRNAs targeting 18,010 genes. Following 2 μg/mL puromycin selection for 72 hours, doxycycline was added for 72 hours to induce Cas9 expression. The cell line used for CRISPRi screen was generated by lentiviral transduction of MCF10A *TP53^-/-^* cells with lenti-BLAST-dCas9-KRAB (Addgene, #89567) followed by selection with 10 μg/mL blasticidin. dCas9-KRAB expressing cells were infected at multiplicity of infection (MOI) 0.3 with a previously published genome-wide human lentiviral CRISPRi library (Horlbeck et al., 2016). The library contains 104,535 sgRNAs targeting 18,905 protein coding genes.

In both CRISPRn and CRISPRi screens, cells were collected for an early time point sample of initial library representation (T_0_) following selection. 10 million CRISPR mutagenized cells were exposed to concentrations that caused a 20% reduction in cell survival (Surviving Fraction 80, SF80) of either olaparib or talazoparib. In total, cells were exposed to drug or DMSO for 14 days (10 population doublings), after which the cells were recovered (T_1_). In order to identify CRISPR guides responsible for modulating PARPi response, sgRNA enrichment and depletion was estimated in cells that survived exposure to PARPi or DMSO using parallel sequencing. In brief, genomic DNA was extracted from T_0_ and T_1_ cells and sgRNA sequences were PCR amplified for sequencing on Illumina sequencing (HiSeq 2500), producing >1,000 short-reads per sgRNA within the library. These short-read sequences were aligned to the known sgRNAs sequences in the respective library, and the frequency of each short-read was determined to estimate sgRNA frequency within the surviving populations.

MAGeCK (Model-based Analysis of Genome-wide CRISPR/Cas9 Knockout) software was used to generate sgRNA counts according to the sequences present in the genome-wide CRISPR library (Li et al., 2014). Using normalised read count data from MAGeCK, quality checks were performed (distribution of read counts, clustering of samples), to confirm the robustness of the data. For downstream analysis of sgRNA read count data, three approaches were used for comparative analysis: (1) MAGeCK (2) z-score and (3) normZ. From MAGeCK workflow, we extracted a ranked list of positively selected hits generated using its robust ranking aggregation algorithm (RRA) approach (Li et al., 2014). For the z-score approach, the low abundant guides with a read count of zero in the T_0_ sample were first identified and removed from the analysis. Then, raw read counts were converted to parts per ten million (pptm) counts to account for variation in the amount of DNA sequenced. The raw pptm counts were log_2_ transformed after adding a pseudo count of 0.5, and the viability effect (VE) z-scores and drug effect (DE) z-scores were calculated. The difference in abundance of sgRNA-specific short reads between drug-treated and DMSO conditions was quantified into DE Z-scores, as previously described (Colic et al., 2019), which quantifies the extent to which an sgRNA construct modulates drug response. This score was corrected to account for viability effects of sgRNA targeting of a specific gene in the absence of drug by calculating the Z-score between DMSO T_0_ and T_1_ samples (VE). DE Z-scores and VE Z-scores were normalised to median absolute deviation (MAD). In order to remove variation in drug effect that can be attributed to VE, a linear model of DE vs VE is plotted, which is used to adjust DE. Ranks for negative selection were generated by sorting results based on their z-score in ascending order. A final list of hits was consolidated from the three approaches by taking the rank product of their ranks.

Increased or decreased representation of sgRNAs in drug samples indicated resistance-causing or sensitisation effect, respectively; positive drug effect Z-scores indicate resistance-causing effects, while negative drug effect Z-scores indicate sensitisation effects. Gene level drug effect Z-scores were calculated to provide an estimate of overall effect size of each gene. In these specific screens, genes were scored as “hits” for sensitisation effects with a viability-correct drug effect Z-score of ≤ –3, with two or more significant sgRNAs. *P* = 0.05 was considered to be statistically significant.

When comparing CRISPR screen results from different investigators, we performed quantile normalisation in order to account for any technical variation across samples. It is a data transformation technique for making two or more data distributions statistically identical to each other. Quantile normalisation was done using the R package, preprocessCore built under R version 4.0.3.

### Retroviral mutagenesis screen

The retroviral mutagenesis screen was performed as described in (Francica et al., 2020) and analyzed as previously described (Blomen et al., 2015). The identified candidates were required to pass an FDR-corrected binominal test with *p*<0.05, an FDR-corrected Fisher’s exact test with *p* <0.05 comparing the IR screens with the four wild-type control screens and had to be either depleted or enriched for sense integrations in both replicates.

### Cell viability and clonogenic assays

Cell viability was quantified using the CellTiter-Glo® assay (Promega) following exposure to various concentrations of drug for five days. Colony formation assay was used to assess long-term drug exposure effects following 10-14 days continuous drug exposure, as described previously (Farmer et al., 2005); drug-containing medium was refreshed every 3 days and cells were stained at assay endpoint with sulforhodamine B. Surviving fraction was calculated for each drug dose relative to DMSO-exposed cells to generate dose/response survival curves.

### Growth assays

Colony formation was estimated using 0.1% crystal violet and colonies were quantified in an automated manner with macros using Image J. Cell viability was quantified using CellTiter-Blue® Cell Viability Assay (#G8081, Promega). Cells were treated with the indicated drug or irradiated at the indicated dosages 24 hours after seeding. IR-treated cells were subsequently exposed to repeated irradiation on day 2 and 3. Olaparib-treated cells were constantly exposed to olaparib during the course of the experiments. Control wells of the 6-well plates were fixed and stained on day 8, whereas treated cells in 6-well plates were fixed and stained on day 11.

### Immunofluorescence

Cells were plated onto coverslips. The following day, cells were fixed either 16 hours post 10 µM olaparib treatment or 3 hours post IR (10 Gy) exposure. Control cells were either exposed to DMSO or no IR. Cells were washed with PBS and permeabilized following fixation. Cells were washed 3 times and blocked for 30 minutes at RT, incubated with the primary antibody for 1 hour at RT with anti-RAD51 (sc-8349 (H-92), Santa Cruz) for MCF10A human cell lines or rabbit-anti-RAD51 (70-001, BioAcademia) for the KB1P-G3B1 mouse cell line. Rabbit-anti-CENPB (#ab259855, Abcam) was also used. Mouse-anti-phospho-H2AX (RRID: AB_309864, Milipore) was used with both human and mouse cell lines. After washing 3 times, cells were incubated with the secondary antibody for 1 hour at RT with Goat anti-Rabbit IgG (H+L) Cross-Adsorbed Secondary Antibody, Texas Red-X (RRID: AB_2556779, #T-6391, Thermo Fisher Scientific) or Goat anti-mouse IgG (H+L) Highly Cross-Adsorbed secondary antibody, Alexa Fluor 488 (RRID: AB_2534088, Thermo Fisher Scientific), washed 3 times, counterstained with DAPI (Life Technologies, 1:50000 dilution) and washed 5 times more before mounting. Z-stack images were acquired using the DeltaVision Elite widefield microscope (GE Healthcare Life Sciences) or Marianas advanced spinning disk confocal microscope (3i). All nuclei were detected by the “analyze particles” command in Fiji. The foci were then quantified with the “find maxima” command.

### Proximity ligation assays (PLAs)

PLAs were performed according to the Duolink Detection Kit protocol (#DUO92101, Sigma Aldrich) with indicated primary antibodies rabbit-anti-HA-Tag (#3724, Cell Signaling) or mouse-anti-HA-Tag (#901501, BioLegend, rabbit-anti-RAD51 (#70-001, Bioacademia), mouse-anti-phospho-H2AX (RRID: AB_309864, Milipore), mouse-anti-Biotin (#200-002-211, Jackson ImmunoResearch). Z-stack images were acquired using the DeltaVision Elite widefield microscope (GE Healthcare Life Sciences). All nuclei were detected by the “analyze particles” command in Fiji. The PLA foci were then quantified with the “find maxima” command. SIRF assay was performed by labelling cells with EdU (25 µM) for 10 minutes followed by three washes with PBS. Replication stress was induced in the HU-treated samples by adding 2 mM HU for 2 hours. Cells were then washed twice with PBS and nuclei were pre-extracted with CSK buffer (10 mM PIPES pH 7, 0.1 M NaCl, 0.3 M sucrose, 3 mM MgCl2, 0.5 % (v/v) Triton X-100) on ice for 5 minutes. After pre-extraction, cells were washed with PBS and fixed in 4% (v/v) PFA/PBS for 20 min on ice. Followed by PLA staining according to manufacturer’s protocol.

### Sister chromatid exchange (SCE) assay

HAP1 cells were treated for 48 hours with 10 µM BrdU. If indicated, 0.5 µM olaparib was added simultaneously with BrdU. Colcemid was added for the last 3 hours at a concentration of 100 ng/mL. Cells were treated with 3 Gy IR 8-10 hours prior to fixation. Hypotonic solution 0.075 M KCl solution was used prior to fixation in 3:1 methanol:acetic acid solution. Metaphase spreads were made by dropping the cell suspensions onto microscope glasses from a height of ∼ 30 cm. Slides were stained with 10 µg/mL bis-Benzimide H 33258 (Sigma) for 30 minutes, exposed to 245 nM UV light for 30 minutes, incubated in 2x SSC buffer (Sigma) at 60°C for 1 hour, and stained in 5% Giemsa (Sigma) for 15 minutes.

### DNA fiber assay

Fork progression was measured as described previously in (Schmid et al., 2018) with a few modifications. Briefly, asynchronously growing subconfluent cells were labeled with 30 μM thymidine analogue 5-chloro-2’-deoxyuridine (CIdU) (#C6891, Sigma-Aldrich) for 20 minutes, washed three times with warm PBS and subsequently exposed to 250 μM of 5-iodo-2′-deoxyuridine (IdU) for 20 minutes. In the experiment assessing RF stability, IdU pulse was followed by adding medium containing 8 mM hydroxyurea (HU) for 6 hours for KB1P-G3B1 cells or 4 mM HU for 3 hours for KB1P-G3 cells. In order to assess RF reversal KB1P-G3 cells, 600nM Mitomycin C (MMC) was used ahead of pulse labeling with CldU and IdU, as described before. MMC treatment was maintained during labeling. All cells were then collected and resuspended in cold PBS at 3.5 x 10^5^ cells/mL density. The labeled cells were mixed 1:5 with unlabeled cells resuspended in cold PBS at 2.5 x 10^5^ cells/mL density. Cells were then resuspended in lysis buffer (200 mM Tris-HCl, pH 7.4, 50 mM EDTA, and 0.5% (v/v) SDS) on a positively-charged microscope slide. After 9 minutes incubation at RT, the DNA fibres were stretched, air-dried, fixed in 3:1 methanol/acetic acid, and stored at 4 °C overnight. The following day, the DNA fibers were denatured by incubation in 2.5 M HCl for 1 hour at RT, washed five times with PBS and blocked with 2% (w/v) BSA in 0.1% (v/v) PBST (PBS and Tween 20) for 40 minutes at RT while gently shaking. The newly replicated CldU and IdU tracks were stained for 2.5 hours at RT using two different anti-BrdU antibodies recognizing CldU (#ab6326, Abcam) and IdU (#347580, Becton Dickinson), respectively. After washing five times with PBST (PBS and Tween-20), the slides were stained with goat-anti-mouse IgG (H+L) Cross-Adsorbed Secondary Antibody, Alexa Fluor 488 (RRID: AB_2534088, #A-11029, Thermo Fisher Scientific) diluted 1:300 in blocking buffer and with the Cy3 AffiniPure F(ab’)₂ Fragment Donkey Anti-Rat IgG (H+L) antibody (#712-165-513, Jackson ImmunoResearch) diluted 1:150 in blocking buffer. Incubation with secondary antibodies was carried out for 1 hour at RT in the dark. The slides were washed five times for 3 minutes in PBST, air-dried and mounted in Fluorescence mounting medium (#S3023, Dako). Fluorescent images were acquired using the DeltaVision Elite widefield microscope (GE Healthcare Life Sciences). To assess RF progression CldU + IdU track lengths of at least 120 fibers per sample were measured using the line tool in ImageJ software. RF stability was analyzed by measuring the track lengths of CldU and IdU separately and by calculating IdU/CldU ratio.

### siRNA transfection

Lipofectamine RNAimax was used according to the manufacturer’s instructions (Invitrogen). Cells were transfected with Smart Pools or deconvoluted siRNA targeting the appropriate gene (GE Dharmacon).

### Western blotting

Standard protocols for SDS–PAGE and immunoblotting were used (Henderson and Wolf, 1992). Proteins were transferred from polyacrylamide gels to nitrocellulose (GE Healthcare) membranes.

### qRT-PCR

Total RNA was isolated (Qiagen RNAeasy) and reverse transcribed with high Capacity cDNA Reverse Transcription Kit (ThermoFisher), as per kit instructions. The RT reactions were amplified with TaqMan probes Hs01552130_g1 MND1 TaqMan probe human (4351372, Thermo), Hs00917175_g1 PSMC3IP TaqMan probe human (4351372, Thermo) and Hs02786624_g1 GAPDH TaqMan probe human (4448489, Thermo) with TaqMan master mix, QuantStudio 6 Flex Real-Time PCR System (Thermo) used for quantification. Fold depletion for each siRNA treatment was determined as 2^ΔΔCt^, for which the cycle threshold (Ct) value for the target mRNA was subtracted by Ct value for GAPDH (mean of duplicate amplifications from the same RT reaction) to calculate the ΔCt value, which was then subtracted from the corresponding ΔCt from siCTRL treated cells to calculate ΔΔCt.

### Cell cycle analysis

HAP1 cells were either left untreated, treated for 16 hours with nocodazole (250 ng/mL), or treated with 2.5 Gy IR (Cesium137 source, CIS Bio international/IBL637 irradiator) at 30 minutes prior to 16 hours nocodazole treatment. Cells were fixed in ice-cold 70% ethanol and stained with propidium iodide (50 μg/mL)/RNase (100 μg/mL) and rabbit-anti-phospho-HistoneH3-antibody (#9701, Cell Signaling) and goat-anti-rabbit-Alexa-488-conjugated secondary antibody (#R37116, Thermo Fisher). At least 10,000 events were analyzed per sample on a Quanteon (Agilent).

### Reagents

Drugs: talazoparib was purchased from Selleckchem and RAD51i B02 was purchased from Sigma. Olaparib was provided by AstraZeneca. Antibodies utilized for Western blotting are as follows: Anti-MND1 (HPA0434, Atlas), anti-PSMC3IP (HPA044439, Atlas), anti-vinculin (sc-73614, Santa Cruz), anti-HA-tag C29F4 (3724, Cell Signalling), anti-actin (A2228, Sigma), anti-actin anti-V5-tag (Cell Signalling, 13202), anti-Cas9 (NBP2-36440, Novus), anti-actin (C-2) (sc-8432, Santa Cruz).

### Statistical analysis

In dose/response survival curves, error bars represent standard deviation (SD) from typically n=3 replicates. In scatterplots, error bars represent the median and 95% confidence intervals (CI). *P*-values were calculated via ANOVA with Tukey’s or Bonferroni post-test.

## Acknowledgements

We thank Jeremy Stark (City of Hope) for the kind gift of U2OS-DR-GFP cells and Thijn Brummelkamp (NKI) for providing HAP1 cells. This work was funded by the Swiss National Science Foundation (310030_179360 to RSR), the European Research Council (CoG-681572 to SSR) the Swiss Cancer League (KLS-4282-08-2017 to SR),the Wilhelm-Sander Foundation (no. 2019.069.1 to SR), by Breast Cancer Now (as part of Programme Funding to CJL and ANJT as part of the Breast Cancer Now Toby Robins Research Centre), by Cancer Research UK (as part of Programme Funding to SJP and CJL). We also thank the Breast Cancer Now Toby Robins Research Centre Bioinformatics Core for Bioinformatics Support and thank Breast Cancer Now, working in partnership with Walk the Walk for supporting the work of this team. We also thank Dr. Kai Betteridge and their team in the ICR Light Microscopy Facility for assistance with microscopy. This work represents independent research supported by the National Institute for Health Research (NIHR) Biomedical Research Centre at The Royal Marsden NHS Foundation Trust and the Institute of Cancer Research, London. The views expressed are those of the author(s) and not necessarily those of the NIHR or the Department of Health and Social Care.

