## Supplementary_figures for "MND1 and PSMC3IP control PARP inhibitor sensitivity in mitotic cells"

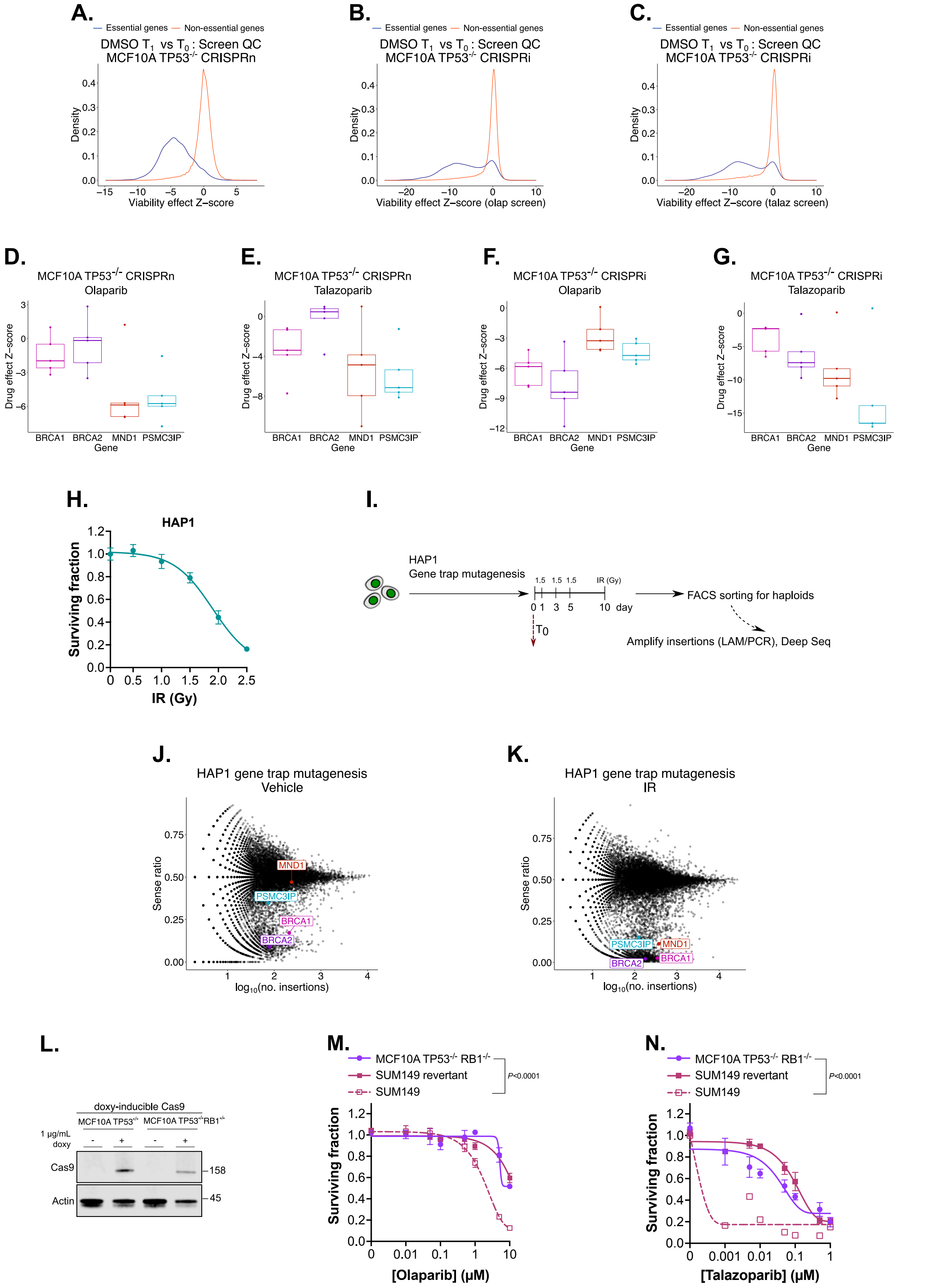


**Supplementary Figure 1. Genetic screens identify MND1 and PSMC3IP as determinants of PARPi and ionizing radiation sensitivity**. **A-C**. Quality of the CRISPRn and CRISPRi screens was assured by depletion of sgRNA targeting essential genes. Scatter plots are shown with viability effect Z-score *vs.* density from genome-wide CRISPRn (A) and

CRISPRi (B, C) screens. The olaparib and talazoparib arms of the CRISPRn (A) screens were performed simultaneously, so the viability effect Z-scores are the same, whereas the olaparib and talazoparib arms of the CRISPRi screens (B, C) were performed at different times. MAGeCK (Model-based Analysis of Genome-wide CRISPR/Cas9 Knockout) software was used to generate sgRNA counts according to the sequences present in the genome-wide CRISPR library (see methods). **D-G.** Box plots of drug effect Z-scores for each sgRNA targeting *MND1* and *PSMC3IP*, from screens described in Figure 1D, E. Effect of MND1 and PSMC3IP is compared to effects elicited via CRISPRn or CRISPRi of *BRCA1* or *BRCA2*. MAGeCK (Model-based Analysis of Genome-wide CRISPR/Cas9 Knockout) software was used to generate sgRNA counts according to the sequences present in the genome-wide CRISPR library (see methods). **H.** Ionising radiation (IR) resistance in HAP1 cells used for retroviral mutagenesis screen outlined in I. Dose/response survival curves are shown with surviving fractions at the indicated doses of IR. Cells were plated in 24-well plates and exposed to indicated dose of IR, after which cell viability was quantified via CellTiter-Blue®. Surviving fraction was calculated for each IR dose relative to cells which remained unexposed to IR. Error bars represent SD from n=6 replicates. **I.** Schematic representing workflow for retroviral mutagenesis screen shown in Supplementary figure 1J, K. **J, K.** Data from retroviral mutagenesis screens comparing control (J) to IR-treated (H). Fishtail plots are shown. Genes with low sense ratio and high log10(no. insertions) represent IR sensitivity-causing effects (as shown by named HR/DNA repair genes). **L.** Western blot image of MCF10A *TP53*$^{-/-}$ cell lysates with or without *RB1* defect illustrating expression of doxycycline-inducible Cas9 (uncropped image shown in Supplementary Figure 12G). **M, N.** PARPi resistance in MCF10A *TP53*$^{-/-}$ cells with *RB1* defect. Dose/response survival curves are shown with surviving fractions at the indicated doses of olaparib (M) or talazoparib (N). Cells were plated in 384 well-plates and exposed to PARPi for five continuous days, after which cell viability was quantified by CellTiter-Glo®. Surviving fraction was calculated for each drug dose relative to DMSO-exposed cells. PARPi sensitive *BRCA1* mutant SUM149 and PARPi resistant *BRCA1* revertant SUM149 cells are shown as controls. Error bars represent SD from n=3 replicates. *P*-values were calculated via ANOVA with Tukey's post-test.

**Supplementary Figure 2.**

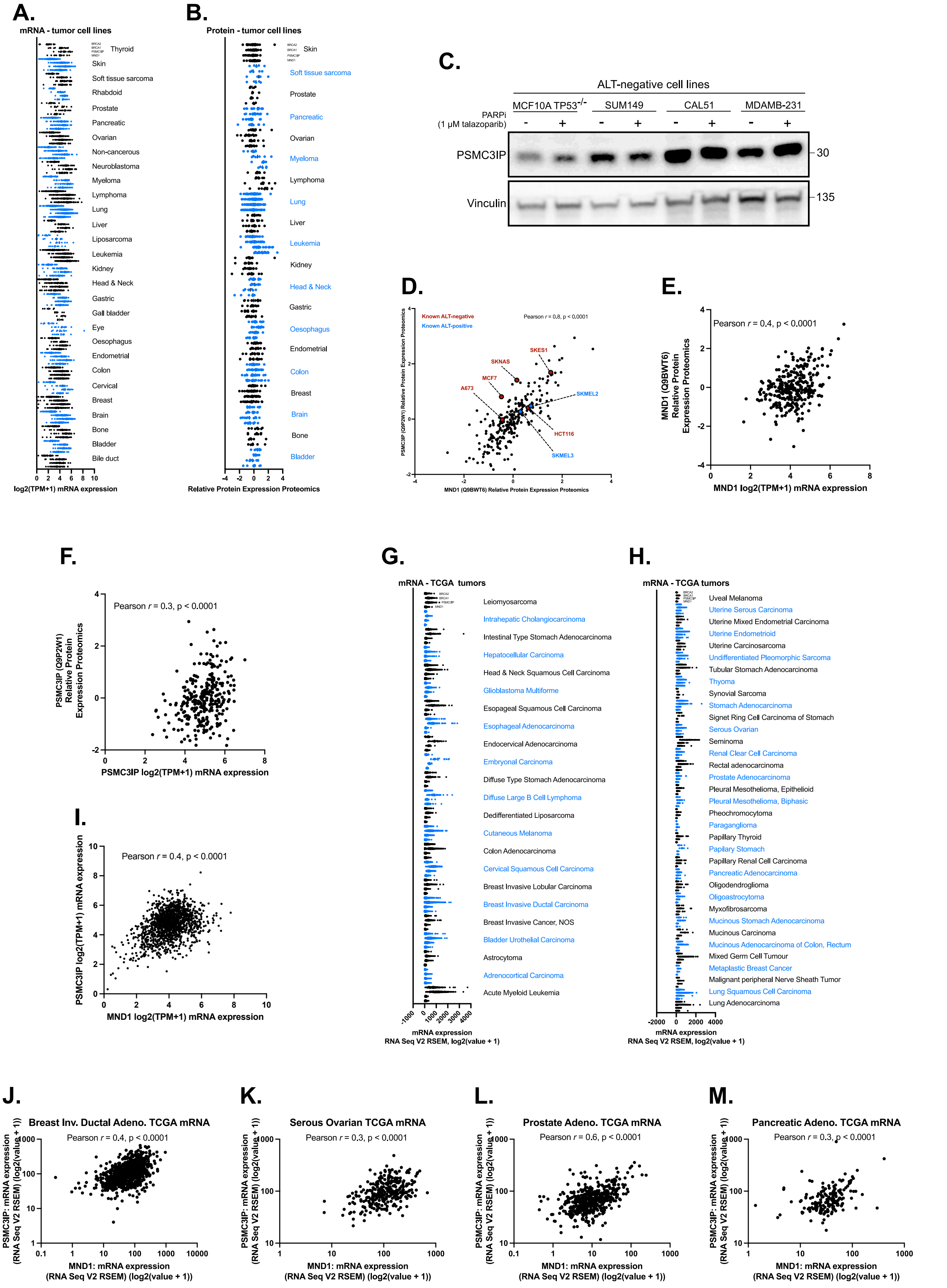


**Supplementary Figure 2. Expression of MND1 and PSMC3IP in mitotic cells.**
**A.** Normalized mRNA expression for *MND1* and *PSMC3IP* in 1407 tumor cell lines profiled as part of the DepMap project (Ghandi et al., 2019). Data for *BRCA1* and *BRCA2* is also

shown as a comparison. Raw data for plot retrieved from **https://depmap.org/portal/** on 1st July 2022 and described in Supplementary Table 11. **B.** Normalized protein expression for MND1 and PSMC3IP in 352 tumor cell lines profiled as part of the DepMap project (Ghandi et al., 2019). Data for *BRCA1* and *BRCA2* is also shown as a comparison. Raw data retrieved from **https://depmap.org/portal/** on 1st July 2022 and described in Supplementary Table 11. **C.** A Western blot image demonstrating PSMC3IP expression in ALT-negative cell lines (Hu et al., 2021). Uncropped image shown in Supplementary Figure 12H. **D, E.** Correlation of PSMC3IP *vs.* MND1 protein expression in tumor cell lines from (A, B). Examples of known ALT-positive and ALT-negative cell lines are highlighted in blue and red, respectively. Data for this plot is listed in Supplementary Table 12, which is adapted from protein expression data shown in Supplementary Table 11 but also includes ALT status of cell lines, if known**.** Correlation data of mRNA *vs.* protein levels for MND1 and PSMC3IP shown from data in (A) and (B) is listed in Supplementary Tables 13 and 14**.** Pearson's correlation coefficient was used. **F, G** Correlation of PSMC3IP *vs.* MND1 mRNA expression in tumor cell lines from (A, B). Data for this plot is listed in Supplementary Tables 11**.** Correlation of mRNA *vs.* protein levels for MND1 and PSMC3IP from data in (A) and (B). Correlation data for these plots is listed in Supplementary Tables 13 and 14**.** Pearson's correlation coefficient used. **H, I.** Normalized mRNA expression for MND1 and PSMC3IP in 10,882 tumors profiled as part of the TCGA project described on the CBio portal (Cerami et al., 2012; Gao et al., 2013)**.** Raw data for plot retrieved from https://www.cbioportal.org/ on 1st July 2022 and described in Supplementary Table 15. **J-M.** Correlation of PSMC3IP *vs.* MND1 protein expression in tumors from four histologies where PARPi are routinely used. Data derived from (H, I) and listed in Supplementary Table 15**.**

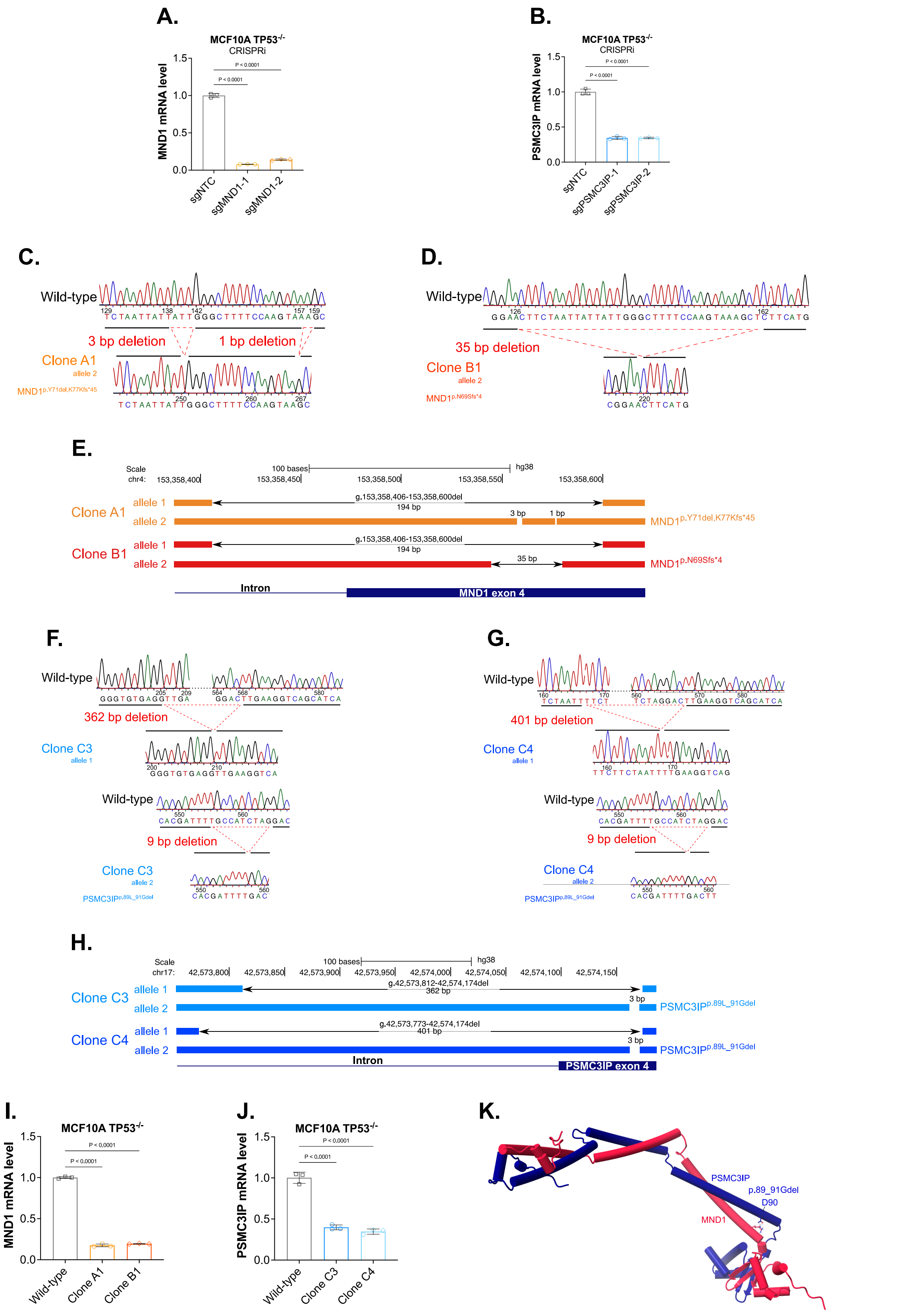


**Supplementary Figure 3. *MND1* and *PSMC3IP* silencing and mutation in mitotic cell lines. A, B.** Depletion of *MND1* (sgMND1) (A) or *PSMC3IP* (sgPSMC3IP) (B) using CRISPRi displayed a significant reduction in either MND1 or PSMC3IP mRNA compared to

MCF10A *TP53*$^{-/-}$ cells expressing non-targeting control (sgNTC). Error bars represent SD from n=3 replicates. *P*-values were calculated via ANOVA with Tukey's post-test. **C, D, E.** Mutations generated in *MND1*-mutant MCF10A *TP53*$^{-/-}$ cell lines. MND1 clones A1 (C, E) and B1 (D, E) had a large genomic deletion g.153,358,406-153,358,600 spanning *MND1* exon 4 and intron in one allele. In addition, clone A1 (C, E) had 3 bp and 1 bp deletions, while clone B1 (D, E) had 35 bp deletion, on the second allele (MND1p.Y71del,K77Kfs*46 and MND1 p.N69Sfs*4, respectively). **F, G, H.** Both PSMC3IP clones C3 (F, H) and C4 (G, H) had a 9 bp deletion p.89L_91G in the second allele. In allele 1, clone C3 (F, H) had a 362 bp deletion g.42,573,812-42,574,174 and C4 (G, H) had a 401 bp deletion g.42,573,773-42,574,174 spanning *PSMC3IP* exon 4 and intron. **I, J.** Mutation of *MND1* (I) or *PSMC3IP* (J) using CRISPR displayed a significant reduction in either MND1 (clone A1 or B1) or PSMC3IP (clone C3 or C4) mRNA compared to unedited MCF10A *TP53*$^{-/-}$ cells (wild-type). Error bars represent SD from n=3 replicates. *P*-values were calculated via ANOVA with Tukey's post-test. **K.** Alfafold2 predicted model of the human PSMC3IP (blue) MND1 (red) heterodimer, with indicated positions of the p.89L_91Gdel mutation site. PSMC3IP D90 is predicted to form a hydrogen bond with Arg82 in MND1.

**Supplementary Figure 4.**

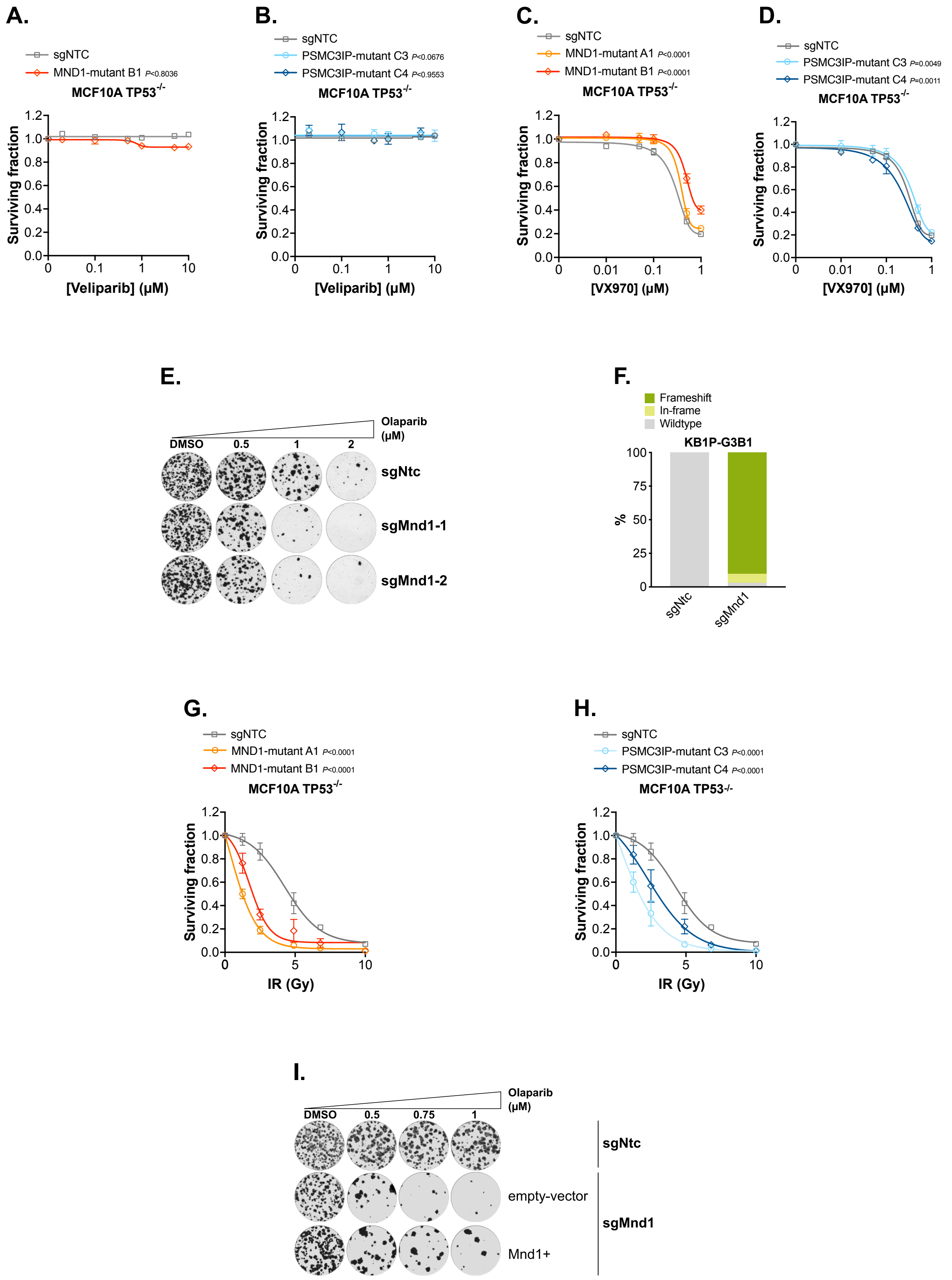


**Supplementary Figure 4. PARPi and IR sensitivity in *MND1* and *PSMC3IP* defective mitotic cells**. **A-D.** Generated *MND1* (A, C) or *PSMC3IP* (B, D) mutant clones were equally resistant to veliparib (A, B) or VX970 (C, D) as unedited cells. Dose/response survival curves are shown with surviving fractions at the indicated doses of veliparib (A, B) or VX970 (C, D). Cells were plated in 384-well plates and exposed to indicated doses of veliparib or VX970 for 7 continuous days, after which cell viability was quantified by CellTiter-Glo®. Surviving fraction was calculated for each drug dose relative to DMSO-exposed cells. Error bars represent SD from n=3 replicates. *P*-values were calculated via ANOVA with Tukey's post-test. **E.** *MND1* defective KB1P-G3B1 cells (sgMnd1) are more sensitive to olaparib than cells expressing non-targeting control (sgNtc). Representative images from Figure 2N are shown. **F.** TIDE analysis demonstrates high *Mnd1* frameshift mutation rate in cells targeted with two different sgRNA against *Mnd1* (sgMnd1-1 or sgMnd1-2) compared to non-targeting control cells (sgNtc). Proportion of cells in grey were determined to be wild-type, those in yellow were determined to have in-frame deletions and those in green were determined to have frameshift mutations. DNA extracted from cells were sequenced via Sanger sequencing and target modifications were confirmed using the TIDE algorithm (Brinkman et al., 2014). **G**, **H.** Generated *MND1* (G) or *PSMC3IP* (H) mutant clones were more sensitive to IR compared to non-targeting control cells (sgNtc). Dose/response survival curves are shown with surviving fractions at the indicated doses of IR. Cells were plated in 96 well-plates and exposed to indicated dose of IR. After 7 days, cell viability was quantified by CellTiter-Glo® and surviving fraction was calculated for each drug dose relative to DMSO-exposed cells. Error bars represent SD from n=8 replicates. *P*-values were calculated via ANOVA with Tukey's post-test. **I.** *Mnd1*-defective KB1P-G3B1 cells (sgMnd1) were more sensitive to IR compared to non-targeting control cells (sgNtc), which was partially reversed with reconstitution of MND1 (Mnd1+). Cells were plated in 6 well-plates and exposed to indicated dose of IR, after which colony formation was estimated by crystal violet staining. Representative images from Figure 2I are shown.

**Supplementary Figure 5.**

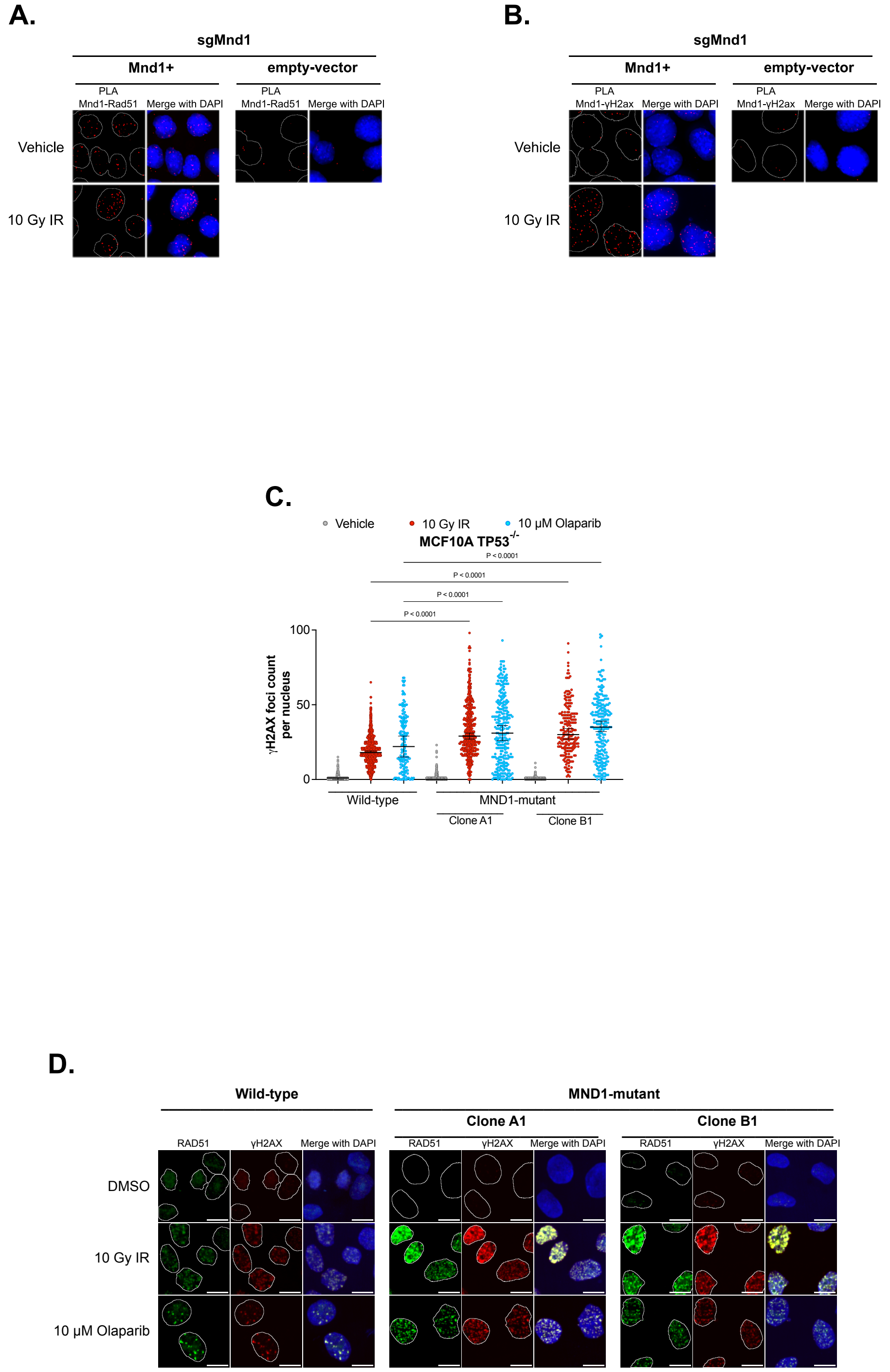


**Supplementary Figure 5. γH2AX and RAD51 foci in cells with *MND1* defects – part 1. A, B.** Representative images of quantification shown in Figures 3A, B demonstrating Mnd1 co-localization with Rad51 (A) in both the presence or absence of exogenous DNA damage, but Rad51 is only co-localized with γH2ax (B) upon exogenous DNA damage. KB1P-G3B1 cells with *Mnd1* defect (sgMnd1), either expressing empty-vector or vector containing Mnd1 cDNA (Mnd1+), were plated on coverslips. Cells were either exposed to 10 Gy IR or remained unexposed. PLAs were performed following staining with anti-HA-tag (tagged to Mnd1) and anti-RAD51 (A) or anti-γH2AX (B) antibodies. **C.** Higher γH2AX foci levels were observed in *MND1* mutant cells compared to wild-type upon olaparib or IR exposure. Scatter plot of RAD51 foci count per nucleus (n= min. 157) in each indicated cell line is shown. MCF10A *TP53*$^{-/-}$ cells, either wild-type or with *MND1* defect (clones A1 and B1) were plated onto coverslips. Cells were either exposed to 10 μM olaparib and then fixed after 16 hours or 10 Gy IR and then fixed after 4 hours, or remained untreated. Cells were co-stained with anti-RAD51 and anti-γH2AX antibodies. Error bars represent the median and 95% CI. *P*-values were calculated via ANOVA with Bonferroni post-test. Representative image shown in D. **D.** Representative images shown of higher RAD51 (Figure 3C) and γH2AX (Supplementary Figure 5C) foci levels observed in *MND1* mutant cells compared to wild-type upon PARPi or irradiation exposure. MCF10A *TP53*$^{-/-}$ cells, either wild-type or with *MND1* defect (clones A1 and B1) were plated onto coverslips. Cells were either exposed to 10 μM olaparib and then fixed after 16 hours or 10 Gy IR and then fixed after 4 hours, or remained untreated. Cells were co-stained with anti-RAD51 and anti-γH2AX antibodies. Scale bar = 10 μm.

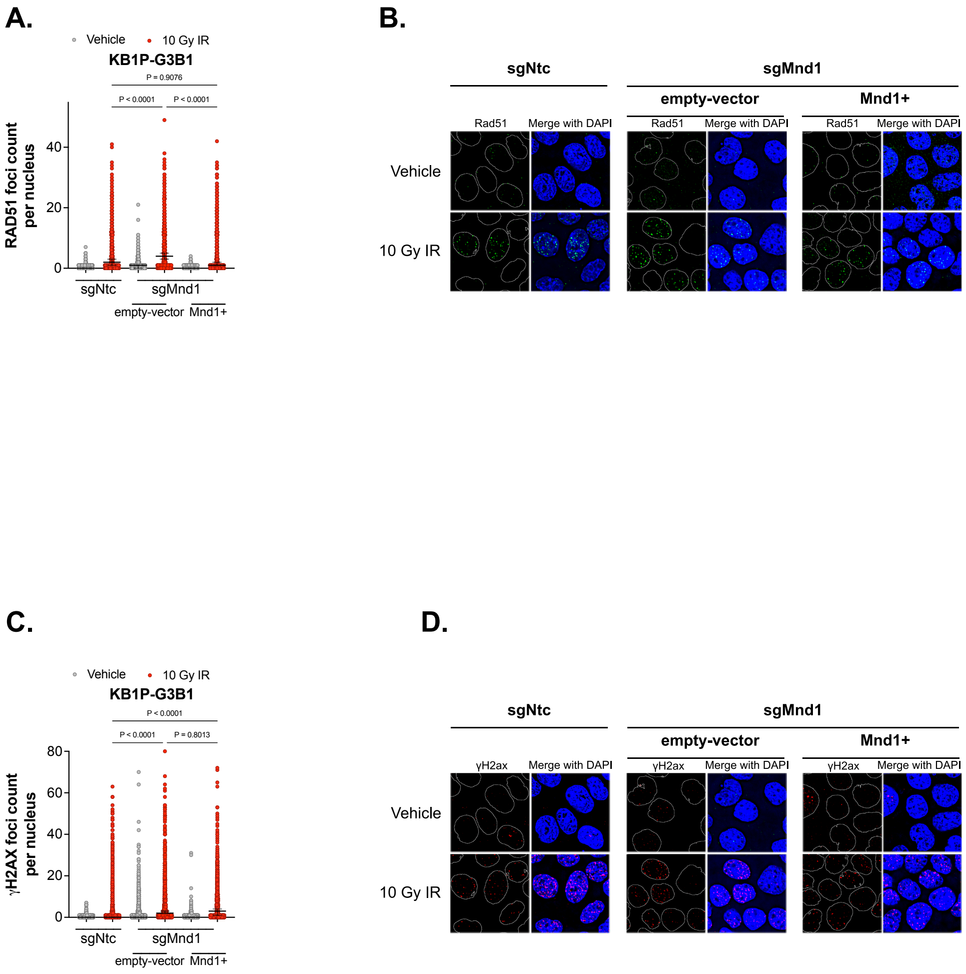


**Supplementary Figure 6. gH2AX and RAD51 foci in cells with *MND1* defects – part 2**. **A, B.** Higher Rad51 foci levels observed in *Mnd1* mutant cells upon IR exposure, which was partially reversed with ectopic Mnd1 expression. Scatter plot of Rad51 foci count per nucleus (n= min. 569) in each indicated cell line is shown. KB1P-G3B1 cells were plated onto coverslips, either expressing nontargeting control or sgRNA targeting *Mnd1* (sgMND1), which expressed either empty-vector or vector containing Mnd1 cDNA (Mnd1+). Cells were either exposed to 10 Gy IR or remained untreated and then fixed. Cells were stained with anti-RAD51. Error bars represent the median and 95% CI. *P*-values were calculated via ANOVA with Tukey's post-test. Representative images from A shown in B. **C, D.** Higher **g**H2AX foci levels observed in *Mnd1* mutant cells upon IR exposure, which was partially reversed with ectopic Mnd1 expression. Scatter plot of **g**H2AX foci count per nucleus (n= min. 580) in each indicated cell line is shown. KB1P-G3B1 cells were plated onto coverslips, either expressing non-targeting control (sgNtc) or sgRNA targeting *Mnd1 (sgMnd1),* which expressed either empty-vector or vector containing *Mnd1* cDNA (Mnd1+). Cells were either exposed to 10 Gy IR or remained untreated and then fixed. Cells were stained with anti-**g**H2AX. Error bars represent the median and 95% CI. *P*-values were calculated via ANOVA with Tukey's post-test. Representative images from C shown in D.

**Supplementary Figure 7.**

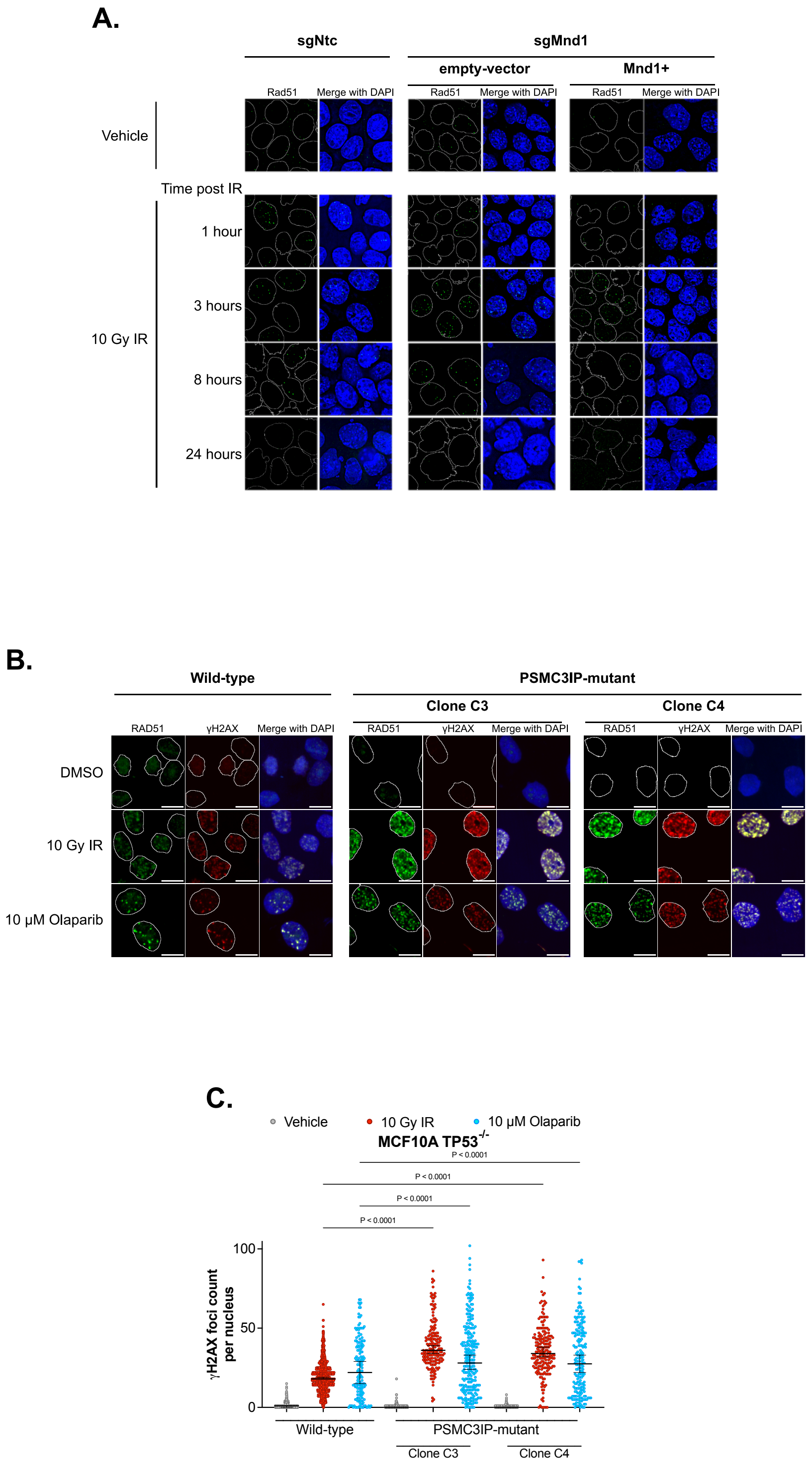


**Supplementary Figure 7. γH2AX and RAD51 foci in cells with *MND1* or *PSMC3IP* defects**. **A.** Increased RAD51 foci levels and decreased kinetics of Rad51 resolution were observed upon IR in *Mnd1* mutant cells. These phenotypes were partially reversed with ectopic Mnd1 expression. Representative images of Rad51 foci in each indicated cell line is shown from quantification in Figure 3D. KB1P-G3B1 cells were plated onto coverslips, either expressing non-targeting control (sgNtc) or sgRNA targeting *Mnd1* (expressing either empty-vector or vector containing Mnd1 cDNA (Mnd1+). Cells were either exposed to 10 Gy IR then fixed at the indicated timepoint or remained unexposed. Cells were stained with anti-RAD51. **B, C.** Higher RAD51 (B) and γH2AX (B, C) foci levels were observed in *PSMC3IP* mutant cells compared to wild-type upon PARPi or IR exposure. MCF10A *TP53*$^{-/-}$ cells were either exposed to 10 μM olaparib and then fixed after 16 hours or 10 Gy IR and then fixed after 4 hours or remained untreated. Cells were co-stained with anti-RAD51 and anti-γH2AX antibodies. Representative images (B) of foci in each indicated cell line is shown from quantification (RAD51 foci in Figure 3E and γH2AX in Supplementary Figure 5C). Scatter plot of γH2AX foci count per nucleus (n= min. 181) in each indicated cell line is shown (C). Error bars represent the median and 95% CI. *P*-values were calculated via ANOVA with Bonferroni post-test.

**A.**

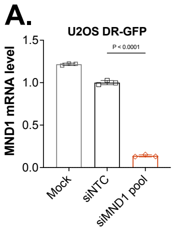


**B.**

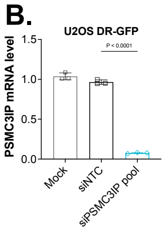


**C.**

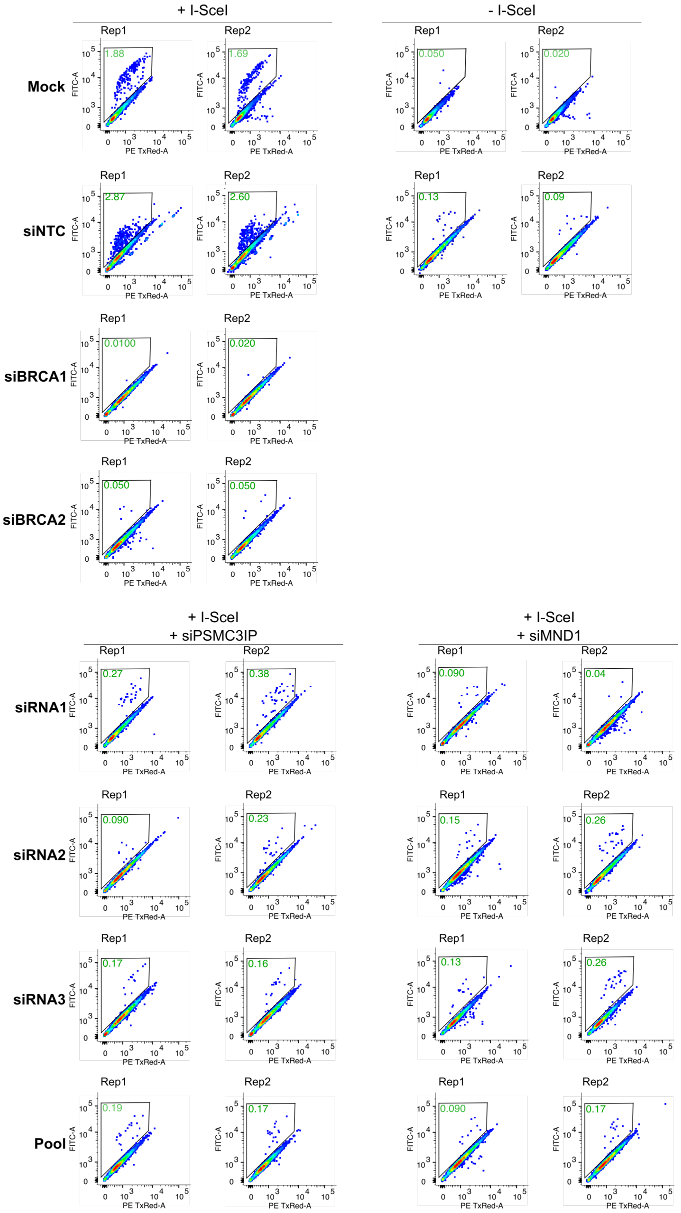


**Supplementary Figure 8. DR-GFP assessment of *MND1* or *PSMC3IP* silenced cells**. **A, B.** Depletion of *MND1* (A) or *PSMC3IP* (B) using siRNA displayed a significant reduction in either MND1 or PSMC3IP mRNA compared to U2OS DR-GFP cells transfected with non-targeting siRNA (siNTC). Normalized to siNTC. Error bars represent SD from n=3 replicates. *P*-values were calculated via ANOVA with Tukey's post-test. **C.** Representative FACS plots of quantification shown in Figure 3F demonstrated *MND1* or *PSMC3IP* silencing reduced HR-mediated repair. GFP+ cells illustrated in green, as analyzed by flow cytometry. U2OS DR-GFP cells (Gunn and Stark, 2012) were treated with siRNAs targeting *MND1, PSMC3IP* or non-targeting control (siNTC), prior to expression of I-*Sce*I. siRNA-mediated silencing of *BRCA1* or *BRCA2* were used as positive controls for HR deficiency.

**Supplementary Figure 9.**

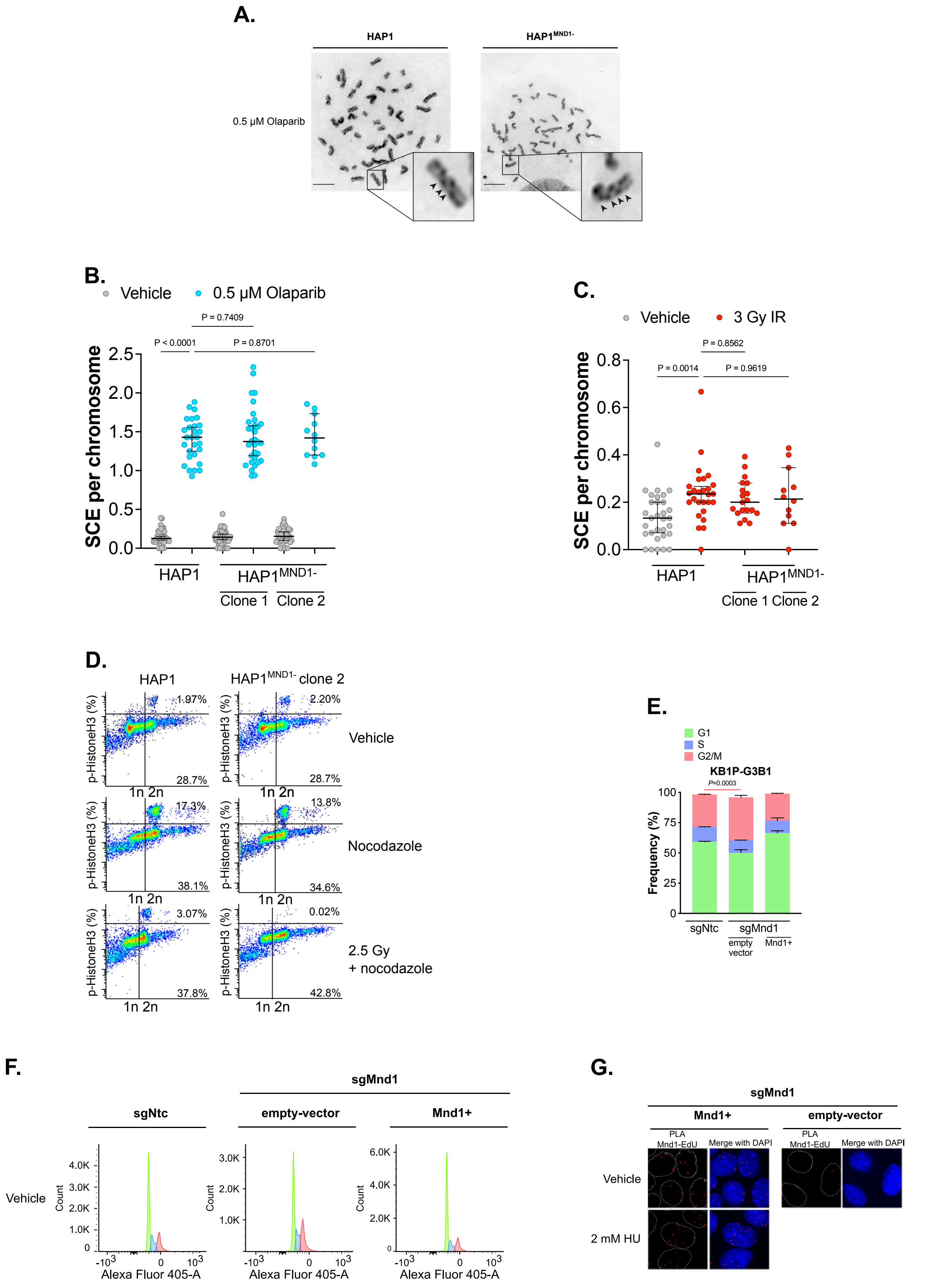


**Supplementary Figure 9. Sister chromatid exchange (SCE) and cell cycle profiling in cells with MND1 or PSMC3IP defects**. **A-C.** *MND1* knockout did not alter the rates of SCEs in untreated (vehicle), IR- or olaparib-treated cells. HAP1 or HAP1$^{MND1-}$ cells were treated with 10 μM BrdU for 48 hours, and 0.5 μM olaparib (B), if indicated. Specified cells were exposed to 3 Gy IR (C) for 8-10 hours. Prior to fixation, cells were treated with hypotonic solution. Metaphase spreads were stained with 10 µg/mL bis-Benzimide H, exposed to UV light and then stained with 5% Giemsa. Representative images demonstrating similar rates of SCE events HAP1 and HAP1$^{MND1-}$ cells upon olaparib-treatment are shown in A. Examples of specific SCE events are highlighted with arrows in blown up images. Scatterplots of SCE per chromosome of olaparib-treated (B) and IR-treated (C) cells are shown. Error bars represent the median and 95% CI from n=3 cells. *P*-values were calculated via ANOVA with Tukey's post-test. **D.** Representative FACS plots from quantification in Figure 3G demonstrate MND1 is required for cell cycle progression following DNA damage. Cells were either left untreated, treated for 16 hours with 250 ng nocodazole per mL, or treated with 2.5 Gy IR at 30 minutes prior to the 16-hour nocodazole treatment. pHistoneH3 was used as a mitotic marker (Wei et al., 1998). **E, F.** A greater proportion of *Mnd1*-defective cells are at $G_2$/M compared to nontargeting control, which is partially reversed with reconstitution of Mnd1 cDNA (Mnd1+). Bar plot of % frequency of cells in specified phase of cell cycle is shown. % cells in $G_1$ phase are shown in green, S phase are shown in purple and $G_2$/M in pink. *P*-values were calculated via t-test. Error bars represent SD from n=3 replicates. Representative FACS plots are shown in F. **G.** Representative images of quantification shown in Figure 3H demonstrating Mnd1 co-localization with EdU-labelled nascent DNA, which is further increased by hydroxy urea (HU)-induced RF stalling. KB1P-G3B1 cells with *Mnd1* defect (sgMnd1), either expressing empty-vector or vector containing Mnd1 cDNA (Mnd1+), were plated on coverslips with EdU. Cells were either exposed to 2 mM hydroxy urea (HU) for two hours or remained unexposed. PLAs were performed following staining with anti-HA-tag (tagged to Mnd1) and anti-biotin antibodies.

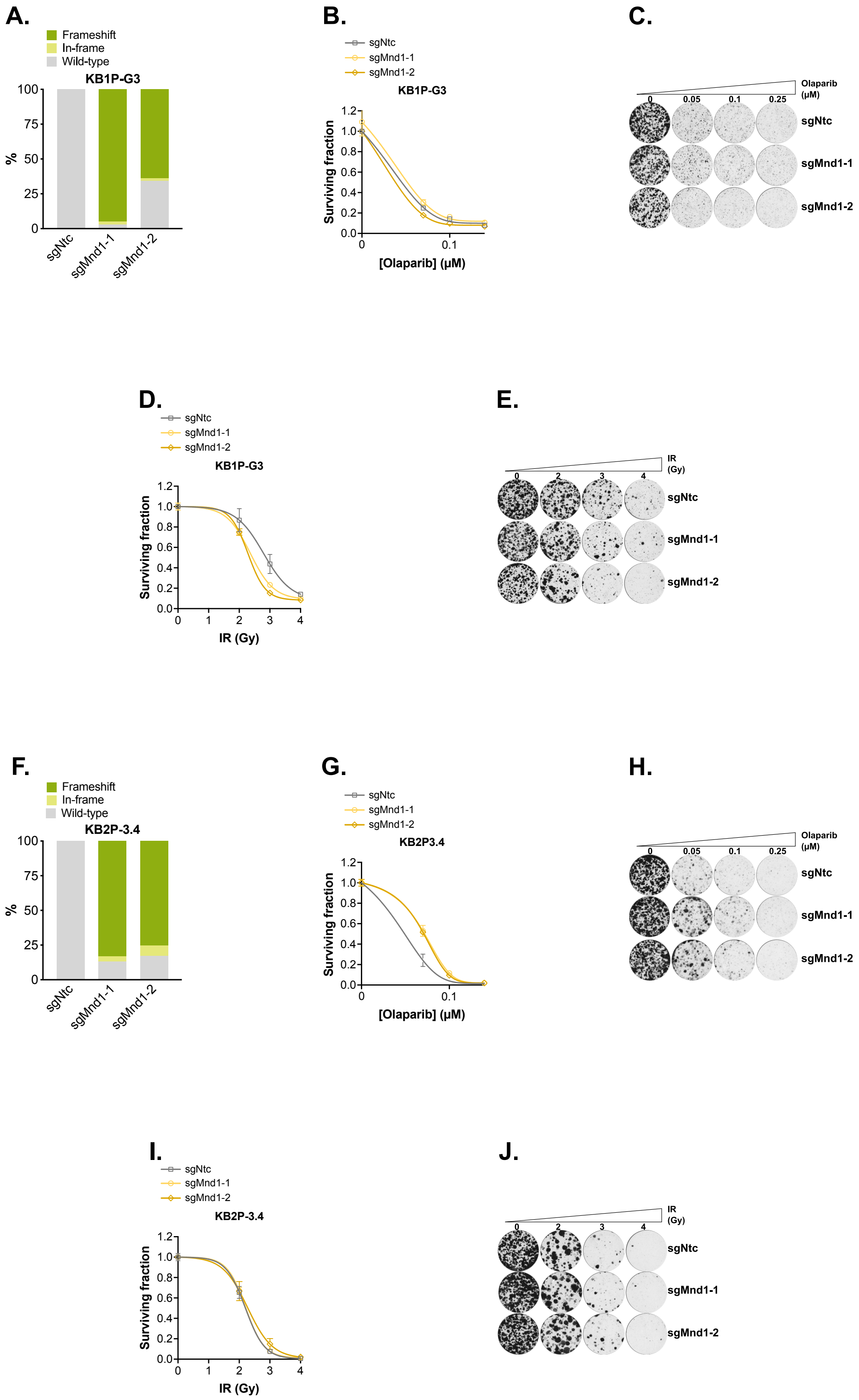


**Supplementary Figure 10. BRCA1/2 epistasis and micronuclei assessment**. **A.** TIDE analysis demonstrates high Mnd1 frameshift mutation rate in KB1P-G3 cells targeted with two different sgRNA against *Mnd1* (sgMnd1-1 or sgMnd1-2) compared to non-targeting control cells (sgNtc). Proportion of cells in grey were determined to be wild-type, those in yellow were determined to have in-frame deletions and those in green were determined to have frameshift mutations. DNA extracted from cells were sequenced via Sanger sequencing and target modifications were confirmed using the TIDE algorithm (Brinkman et al., 2014). **B-E.** *Mnd1* is epistatic with defects in *Brca1.* Compared to control cells, we did not observe further olaparib (B, C) or IR (D, E) sensitization with *Mnd1* defect in *Brca1*-mutant KB1P-G3 cell lines. Cells were plated in 6-well plates and exposed to indicated dose of olaparib (B, C) or IR (D, E) and then cultured for 11 continuous days, after which colonies were stained with crystal violet; colonies were quantified in an automated manner with macros using ImageJ. Surviving fraction was calculated for each drug dose relative to DMSO-exposed cells. Dose/response survival curves are shown in B, D and representative images in C, E, respectively. Error bars represent SD from n=2 replicates. *P*-values were calculated via ANOVA with Tukey's post-test. **F.** TIDE analysis demonstrates high Mnd1 frameshift mutation rate in KB2P-3.4 cells targeted with two different sgRNA against *Mnd1* (sgMnd1-1 or sgMnd1-2) compared to non-targeting control cells (sgNtc). Proportion of cells in grey were determined to be wild-type, those in yellow were determined to have in-frame deletions and those in green were determined to have frameshift mutations. DNA extracted from cells were sequenced via Sanger sequencing and target modifications were confirmed using the TIDE algorithm (Brinkman et al., 2014). **G-J.** *Mnd1* is epistatic with defects in *Brca2.* Compared to control cells, we did not observe further olaparib (G, H) or IR (I, J) sensitization with *Mnd1* defect in *Brca2*-mutant KB2P-3.4 cell lines. Cells were plated in 6-well plates and exposed to indicated dose of olaparib (G, H) or IR (I, J) and then cultured for 11 continuous days, after which colonies were stained with crystal violet; colonies were quantified in an automated manner with macros using ImageJ. Surviving fraction was calculated for each drug dose relative to DMSO-exposed cells. Dose/response survival curves are shown in G, I and representative images in H, J, respectively. Error bars represent SD from n=2 replicates. *P*-values were calculated via ANOVA with Tukey's post-test. **K, L.** *Psmc3ip* loss increases micronuclei formation upon exposure to olaparib (K) or IR (L). Scatterplots of % cells with micronuclei in each indicated sample are shown (n=20). KB1P-G3B1 expressing either sgRNA targeting *Psmc3ip* or non-targeting (sgNtc) control were plated onto coverslips. Cells were either exposed to 10 μM olaparib for 16 hours or 10 Gy IR, or remained untreated. Error bars represent the median and 95% CI. *P*-values were calculated via ANOVA with Tukey's post-test. Representative images shown in Supplementary Figure R. **M-R,** Increased micronuclei formation was observed in *Mnd1*- (M-O) or *Psmc3ip*-deficient (P-R) KB1P-G3B1 cells exposed to olaparib or IR represent broken chromosomes parts and not missegregation of whole chromosomes, as they were negative for the centromere marker CENP-B. Scatterplots of % cells with micronuclei in each indicated sample are shown (n=20). KB1P-G3B1 expressing either nontargeting control or sgRNA targeting *Mnd1* or *Psmc3ip* were plated onto coverslips. *Mnd1*-deficient cells either express an empty-vector or a vector containing Mnd1 cDNA (Mnd1+). Cells were either exposed to 10 μM olaparib for 16 hours, 10 Gy IR, or remained untreated. DNA was stained with DAPI. Error bars represent the median and 95% CI. *P*-values were calculated via ANOVA with Tukey's post-test. Representative images for quantification of micronuclei in *Mnd1*-mutant (sgMnd1) KB1P-G3B1 cells compared to non-targeting control (sgNtc) in M and N shown in O. Representative images for quantification of micronuclei in *Psmc3ip*-mutant (sgPsmc3ip) KB1P-G3B1 cells compared to non-targeting control (sgNtc) in P and Q shown in R.

**Supplementary Figure 11.**

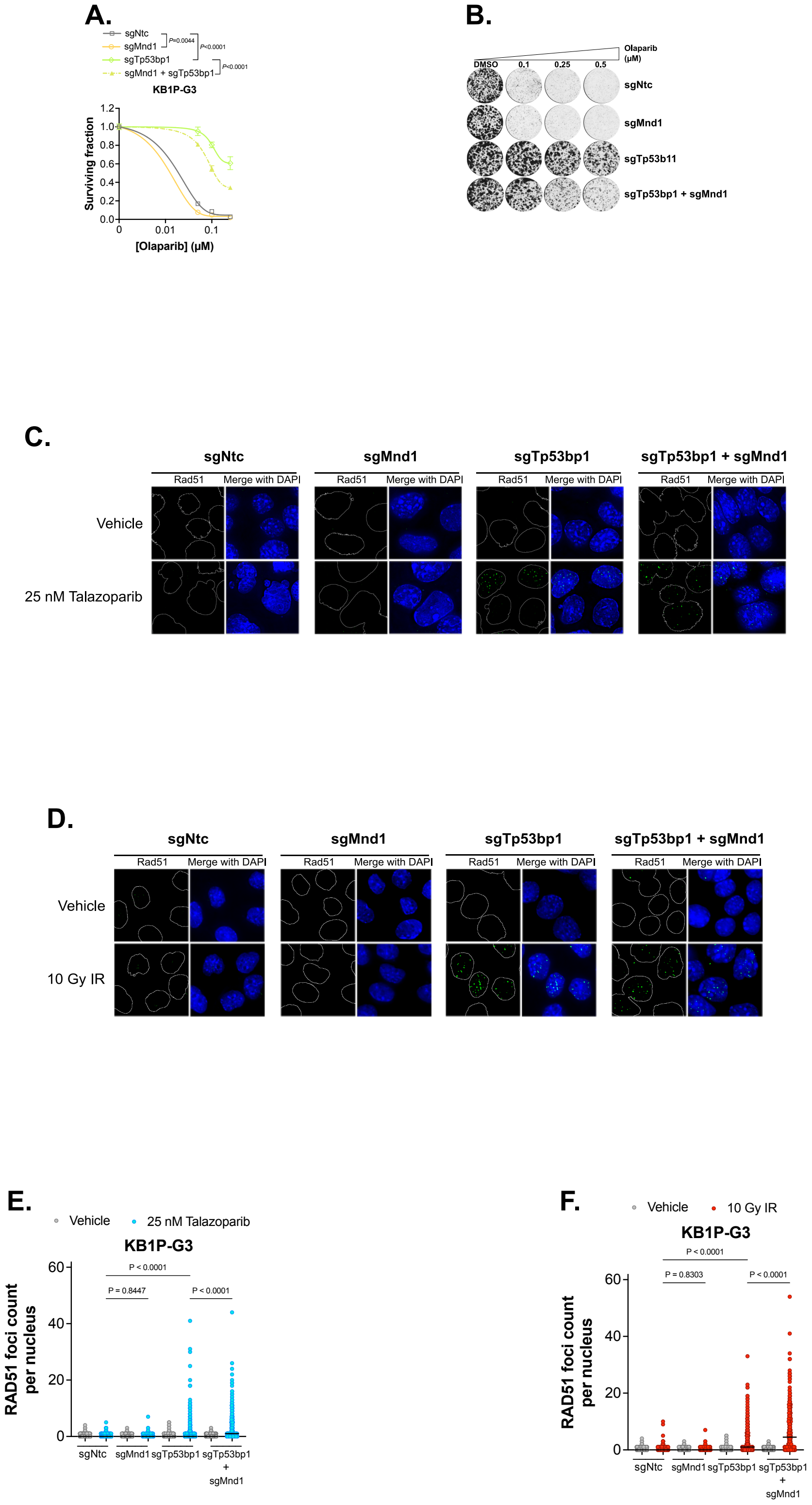


**Supplementary Figure 11. Loss of *Tp53bp1* reverses PARPi sensitivity and increased Rad51 foci in *Mnd1* defective cells. A, B.** *Mnd1* depletion sensitizes *Brca1*- Tp53bp-deficient cells that acquired PARPi resistance by loss of *Tp53bp1*. Dose/response survival curves are shown in A and images of representative colony formation are shown in B. KB1P-

G3 *Brca1*- *Tp53bp1*-deficient cells were plated into 6-well plates, either expressing non-targeting control (sgNtc) or sgRNA targeting *Mnd1 (sgMnd1) or Tp53bp1* (sgTp53bp1) or both *Mnd1* and *Tp53bp1* simultaneously (sgMnd1 + sgTp53bp1). Cells were exposed to olaparib for 11 continuous days, after which colony number was assessed by crystal violet staining; colonies were quantified in an automated manner with macros using ImageJ. Surviving fraction was calculated for each drug dose relative to DMSO-exposed cells. Error bars represent SD from n=2 replicates. *P*-values were calculated via ANOVA with Tukey's post-test. **C-F.** *Mnd1* depletion increased Rad51 foci levels in *Brca1*- *Tp53bp1*-deficient cells that acquired resistance to PARPi talazoparib (C, E) or IR (D, F), by loss of *Tp53bp1.* KB1P-G3 *Brca1*- *Tp53bp1*-deficient cells were plated into 6-well plates, either expressing non-targeting control (sgNtc) or sgRNA targeting *Mnd1 (sgMnd1) or Tp53bp1* (sgTp53bp1) or both *Mnd1* and *Tp53bp1* simultaneously (sgMnd1 + sgTp53bp1). Cells were either exposed to 25 nM Talazoparib (C, E) or 10 Gy IR (D,F) or remained unexposed. Cells were stained with anti-RAD51. Scatter plot of RAD51 foci count per nucleus (n=min 284) in each indicated cell line is shown in E, F. Error bars represent the median and 95% CI. *P*-values were calculated via ANOVA with Bonferroni post-test. Representative images for talazoparib- (C) treated cells quantified in E and IR- (D) treated cells quantified in F.
